# Decorin suppresses tumor lymphangiogenesis: A mechanism to curtail cancer progression

**DOI:** 10.1101/2023.08.28.555187

**Authors:** Dipon K. Mondal, Christopher Xie, Simone Buraschi, Renato V. Iozzo

**Affiliations:** Department of Pathology and Genomic Medicine, and the Translational Cellular Oncology Program, Sidney Kimmel Cancer Center, Sidney Kimmel Medical College at Thomas Jefferson University, Philadelphia, PA 19107, USA

## Abstract

The complex interplay between malignant cells and the cellular and molecular components of the tumor stroma is a key aspect of cancer growth and development. These tumor-host interactions are often affected by soluble bioactive molecules such as proteoglycans. Decorin, an archetypical small leucine-rich proteoglycan primarily expressed by stromal cells, affects cancer growth in its soluble form by interacting with several receptor tyrosine kinases (RTK). Overall, decorin leads to a context-dependent and protracted cessation of oncogenic RTK activity by attenuating their ability to drive a pro-survival program and to sustain a pro-angiogenic network. Through an unbiased transcriptomic analysis using deep RNAseq, we discovered that decorin downregulated a cluster of tumor-associated genes involved in lymphatic vessel development when systemically delivered to mice harboring breast carcinoma allografts. We found that Lyve1 and Podoplanin, two established markers of lymphatic vessels, were markedly suppressed at both the mRNA and protein levels and this suppression correlated with a significant reduction in tumor lymphatic vessels. We further discovered that soluble decorin, but not its homologous proteoglycan biglycan, inhibited lymphatic vessel sprouting in an *ex vivo* 3D model of lymphangiogenesis. Mechanistically, we found that decorin interacted with VEGFR3, the main lymphatic RTK, and its activity was required for the decorin-mediated block of lymphangiogenesis. Finally, we discovered that Lyve1 was in part degraded via decorin-evoked autophagy in a nutrient- and energy-independent manner. These findings implicate decorin as a new biological factor with anti-lymphangiogenic activity and provide a potential therapeutic agent for curtailing breast cancer growth and metastasis.

## INTRODUCTION

The lymphatic system is a unidirectional conduit that recycles protein-rich lymph in the interstitial space back to the venous system and regulates antigen transport to lymph nodes^1;2^. Lymphangiogenesis is a process where new lymphatic vessels are developed from pre-existing lymphatics, a method similar to angiogenesis with blood vessels^3^. During embryonic development, lymphangiogenesis acts as a dynamic process and, importantly, this process is generally inactive in adults under normal physiological conditions. Indeed, adult lymphangiogenesis only occurs in certain pathological conditions such as inflammation and wound healing^1;2^.Unfortunately, lymphatic vessels (LV) also function as a major pipeline for metastasizing cancer cells. Cellular elements from the cancer proper and the tumor stroma produce lymphangiogenic factors such as various vascular endothelial growth factors (VEGF) that facilitate the process of LV sprouting from native LVs^4^. It is well-established that VEGFC can directly bind to the VEGF receptor-3 (VEGFR3) on lymphatic endothelial cells to stimulate lymphangiogenesis^5^.

Formation of new LVs allows tumor cells to migrate by providing smooth channels through which tumor cells can invade and proliferate in secondary loci^6^. Preclinical and clinical evidence suggest that the most common pathway of initial tumor metastasis is through the lymphatic system^4^. Indeed, in many cancers, the detection of tumor metastases in the tumor-draining lymph nodes is considered the first step of tumor dissemination, and recently it has been shown that lymph node colonization induces tumor-immune tolerance to promote distant metastasis^7^. Targeting of the VEGFC/VEGFR3 axis is considered one of the best potential therapeutic strategies against skin cancer metastasis, tumor associated lymphangiogenesis, and lymphatic metastasis^8;9^. Notably, blockade of VEGFC-dependent signaling inhibits lymphatic malformations, both macrocystic and microcystic, driven by the constitutive activation of the p110α PI3K^10^. Thus, a better understanding of lymphangiogenesis is of utmost importance in furthering our knowledge concerning the role of lymphatic vasculature in both primary and metastatic tumors and genetic diseases affecting postnatal lymphangiogenesis.

A key factor in both cancer growth and development comes from the interactions between malignant cells and the molecular and cellular components of the tumor stroma. Bioactive molecules such as proteoglycans often act as interlocutors of this complex interplay, dictating the ability of tumor cells to drive metastatic activity^11^. One of the best studied proteoglycans in the context of cancer growth is the stromal derived decorin, a small leucine-rich proteoglycan (SLRP) with a vast résumé of outside-in signaling^12–15^. Soluble decorin acts as a pan-receptor tyrosine kinase (RTK) inhibitor by suppressing pathways leveraged by tumor cells for growth and survival. We discovered that decorin is a biological ligand for EGFR^16–19^, Met^20;21^, IGF-IR^22^ and VEGFR2^23^, thereby targeting both the cancer proper and the microenvironment. By interacting with EGFR, decorin causes sustained downregulation of the receptor^18^ and degradation via caveosomes^24^ leading to p21*^WAF^*^1^ induction, cell cycle arrest^25;26^, and concurrent upregulation of the antiangiogenic TSP1^27^. Moreover, systemic delivery of recombinant decorin significantly reduces primary tumor growth and eliminates metastases in a highly-metastatic orthotopic breast cancer model^28^. Corroborating evidence showed that treatment with Adenovirus-carrying decorin (Ad.Dcn) causes a great reduction in pulmonary metastases^28–30^. Further systemic Ad.Dcn inhibits skeletal metastases of prostate cancer^31^ and reduces the growth of carcinoma xenografts and contralateral tumors, thus demonstrating a distant antitumor effect^32^. These findings have been independently confirmed by several studies ^33–35^. Moreover, *de novo* expression of decorin inhibits tumor growth and lung metastases of inflammatory breast cancer^36^, and the transcription factor MEIS, a HOXB-binding tumor suppressor, mediates its anti-oncogenic activity by suppressing decorin transcription^37^.

While performing an unbiased transcriptomic analysis using deep RNAseq on mice harboring orthotopic triple-negative breast carcinoma allografts, we serendipitously discovered that systemic decorin delivery downregulated a cluster of tumor-associated genes involved in LV development. Two established lymphatic vessel markers, Lyve1 and Podoplanin, were suppressed at both the mRNA and protein levels *in vivo* correlating with a reduction in tumor volumes. Additionally, we found that decorin, but not its homologous SLRP biglycan, inhibited lymphatic vessel sprouting in an *ex vivo* model of lymphangiogenesis utilizing thoracic duct explants embedded in a 3D collagen matrix. Mechanistically, we found that decorin interacted with VEGFR3, one of the major lymphatic RTKs. Moreover, we found that VEGFR3 activity was required for the decorin-mediated block of lymphangiogenesis and discovered that Lyve1 was in part degraded via decorin-evoked autophagy in a nutrient- and energy-independent manner. Collectively, our findings implicate decorin proteoglycan as one of the few biological factors with anti-lymphangiogenic activity, potentially opening new areas of research based on protein-based therapy to curtail LV growth and development in primary and metastatic solid tumors.

## RESULTS

### Decorin is a Stromal Proteoglycan and its Higher Expression is a Marker of Better Prognosis in Breast Cancer Patients

To assess the role of decorin in human cancer, we queried the Cancer Genome Atlas (TCGA) encompassing >20,000 primary cancers and matched normal samples^38^. We found a significant reduction of *DCN* expression in 18 solid tumors (not shown), especially in breast cancer (*P*<0.001,Fig. 1*A*). Analysis of single-cell RNAseq from a breast cancer cohort^39^ showed that *DCN* is exclusively expressed by immunomodulatory and myofibroblastic cancer associated fibroblasts (CAFs) (Fig. 1*B*), in direct support of our previous studies showing that decorin is a stromal proteoglycan^26;28;40–42^. We corroborated these RNAseq data using the Human Protein Atlas: decorin was specifically detected in the stroma of ductal and lobular breast carcinomas (Fig. 1C). Next, we queried datasets from three different breast cancer cohorts^43–45^ where expression and grade information was curated using ShinyGEO^46^. Notably, *DCN* expression was inversely associated with tumor grade across all cohorts with a significantly lower abundance of *DCN* mRNA in the most aggressive, grade 3 tumors (Fig. 1*D*). Using KMplotter^47;48^ from GEO & EGA repositories, we found that high *DCN* levels correlated with a better relapse-free survival and were a predictive factor independent of clinical stage by multivariate logistic regression analysis (Fig. 1*E*). These data are reinforced by reports where reduced *DCN* expression is associated with poor outcome^49;50^ whereas high stromal *DCN* predicts a better prognosis in breast cancer patients^51^. The anti-oncogenic properties of decorin have also been confirmed in various preclinical^28;29;31;32;35;49;52–58^ and genetic studies utilizing *Dcn^-/-^* mice^12–15;59;60^. This large body of work provides a strong rationale for investigating the mechanism of action whereby a soluble, stromal-derived proteoglycan inhibits breast cancer.

**Fig. 1.**
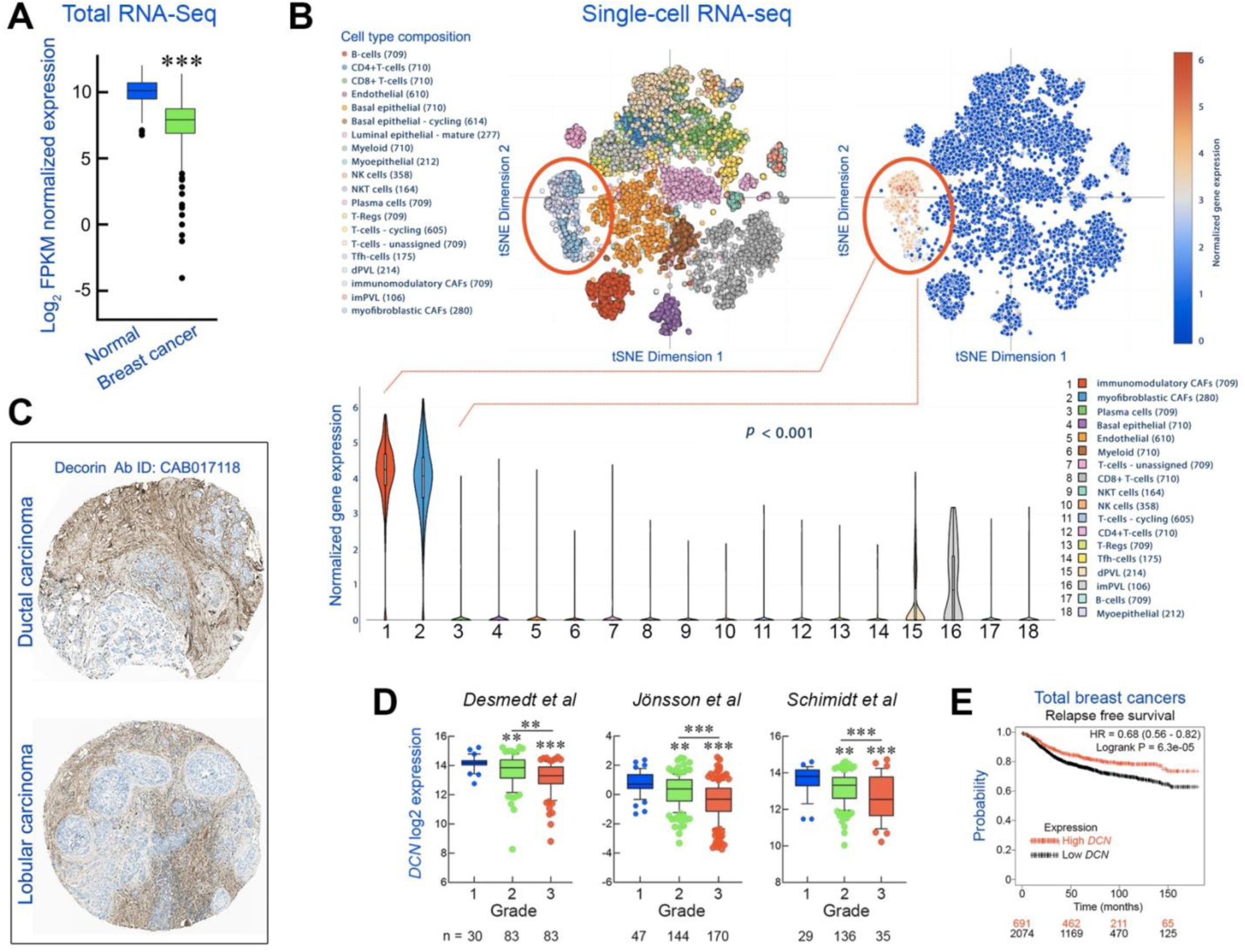
Role of decorin in breast cancer. (*A*) TCGA RNAseq of DCN expression in normal (n=806) and malignant (n=1075) breast carcinoma tissues. (*B*) Single-cell RNAseq of breast cancer patients. (*C*) Stromal DCN expression in ductal & lobular carcinomas. (*D*) DCN expression in 3 different breast cancer datasets stratified by grade; below: samples/grade;\*\**P*<0.01,\*\*\**P*<0.001. (*E*) Relapse-free survival in breast cancers (n= 2765) vis-à-vis DCN expression. The numbers below are colour-coded for high and low DCN expression.

### Systemic Delivery of Recombinant Decorin Suppresses Breast Cancer Growth and Evokes an RNA Signature Including Markers of Lymphatic Vessels

To further explore decorin oncosuppressive activity, we generated an orthotopic breast carcinoma allograft using E0771, triple-negative breast cancer cells derived from C57BL/6 mice^61–64^. These cells have a propensity for spontaneous lung^65^ and bone^66^ metastases, desmoplasia^67^ and are syngeneic to our *Dcn^-/-^* mice^68^, which we backcrossed into the C57BL/6 background for >12 passages. For proteoglycan-based therapy, we purified recombinant His6-tagged decorin expressed in 293-EBNA cells using a Celligen- Plus bioreactor followed by Ni-NTA-affinity chromatography^69^. The samples were assessed for purity via Colloidal Coomassie Blue staining, which has a detection threshold of ∼5 ng^70^. As no additional bands were detected at 1000-fold excess (5 μg, Fig. 2*A*), our samples are >99.9% pure (<5 ng/5000 ng). We treated mice (i.p.,5 mg/kg) starting on day 10 when the tumors became palpable, with subsequent injections every other day until Day 22 (Fig. 2*B*). We found a marked inhibition of growth (*P*<0.001 at Day 22) and established that exogenous decorin indeed targeted the allografts which showed anti-His immunoreactivity in the tumor proper and vessels (Fig. 2*C*). We found a marked reduction of the relative fluorescence of both β-catenin, a known target of decorin^21^, and the endothelial marker CD31^71;72^ (Fig. 2*D*-*F*). We note that we used an unbiased method for quantification: the images were first captured in the vehicle-treated samples and then the objective was moved to the adjacent sections, maintaining constant exposure time, gain and intensity. Finally, we validated these data via WB showing that systemic decorin delivery significantly downregulated an oncogenic (β-catenin) and an angiogenic (CD31) pathway (Fig. 2*G*,*H*).

**Fig. 2.**
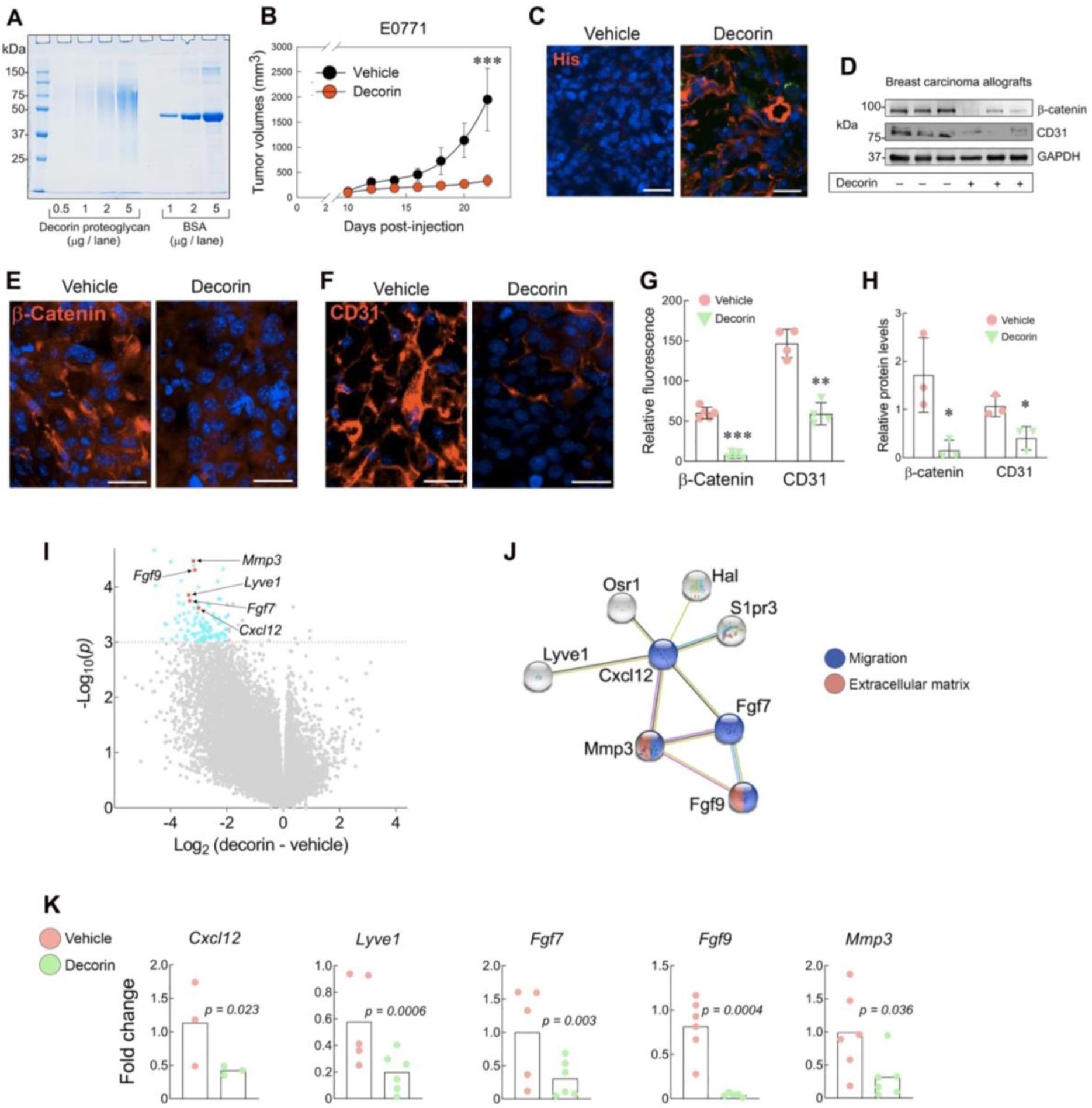
Effects of decorin on E0771 breast carcinoma allografts. (A) SDS-PAGE of endotoxin-free decorin and BSA stained with Colloidal Coomassie Blue. (B) Tumor growth following i.p. injections of PBS or 5 mg/kg DecorinHis6.;***P<0.001, n=7 each. (C) Immunofluorescence images of frozen sections from vehicle or decorin-treated allografts reacted with anti-His antibody. (D,E) Immunofluorescence images of vehicle and decorin treated tumors immunostained for β-catenin or CD31. (F) Quantification of relative fluorescence; n=5, 10 fields/case; Bars ∼ 50 μm.(G) WB of β-catenin and CD31 and (H) Relative quantification, n= 4, *P<0.05). (I) Volcano plot with 5 highlighted genes downregulated >3 fold, P<0.001 adjusted for multiple comparisons; n=7 each for vehicle and decorin. (J) STRING analysis of a subset of RNA-encoded proteins as interaction network with a minimum score of 0.4 (medium confidence). The connecting lines between protein nodes indicate evidence of text mining (green), co-expression (black), and known interactions from databases (blue) or experiments (pink). Nodes are coloured according to indicated GO terms. (K) Validation by qPCR of frozen tumors ±decorin encompassing the 5 downregulated genes highlighted in red.

To gain insight into decorin oncosuppressive effects on the tumor transcriptome, we performed high-throughput RNAseq (Genewiz, n=14 allografts) at day 22 post-treatment (RNAseq data available at www.iozzolab.com). mRNA was enriched and sequenced using Illumina® HiSeq® configured at 2x150 bp. Sequence reads were trimmed to remove possible adapter sequences and poor-quality nucleotides using Trimmomatic v.0.3681 and mapped to the ENSEMBL *Mus musculus* GRCm38 reference genome ENSEMBL using STAR aligner v.2.5.2b^73^. We counted only exonic reads using a strand-specific library and we used the gene hit counts table for differential expression analysis using DESeq2, with the Wald test to generate *p* values and Log2 fold changes. We discovered remarkable changes in the transcriptome of decorin-treated allografts with the vast majority of the genes downregulated (Fig. 2*I*). Gene ontology showed that most of the downregulated genes belonged to ECM and migration. To narrow our search, we applied very stringent criteria for candidate selection (e.g. >2 fold decrease with *P*<0.001). Among the 124 downregulated genes, a cluster of very interesting targets emerged, including *Cxcl12*, *Fgf7/9*, *Lyve1*, and *Mmp3* (Fig. 2*I*). STRING analysis using v11 database 80 revealed that the five-gene signature emerged as interacting proteins, based on predicted or experimentally-proven protein-protein interactions (Fig. 2*J*). We further validated the RNAseq data via qPCR (Fig. 2*K*)). Notably, identification of Fgf7/9 independently supports our previous data on decorin-evoked antiangiogenesis^74–79^.The unexpected suppression of *Lyve1* suggested that decorin could be a novel lymphangiostatic factor.

### Decorin, but not its Homolog Biglycan, Curtails Breast Cancer Growth and Suppresses Lyve1, Podoplanin and Has2 Protein Levels

To further explore the role of decorin in modulating breast cancer development, we performed additional *in vivo* experiments utilizing E0771 breast carcinoma allografts. We confirmed that recombinant decorin significantly inhibited cancer growth in contrast to recombinant biglycan (*P*<0.05, n=7-9 mice/experiment, Fig. 3*A*), the most homologous SLRP to decorin^80^. One of the genes suppressed by decorin in our RNAseq data (cfr. Fig 2) was *Lyve1*, a type I transmembrane receptor homologous to the ubiquitous hyaluronan (HA) receptor CD44^81–83^, and mostly restricted to LVs making it a useful marker for studying tumor LVs^84;85^. To corroborate the RNAseq data, we used specific antibodies against mouse CD31 and Lyve1 and reliably identified vascular and LVs, respectively, in lung, liver and kidney (Fig. 3*B*). We found a marked suppression of Lyve1 in the treated allografts, especially in the intra-tumor LVs (Fig. 3*C*) and a significant reduction of Lyve1 fluorescence (*P<*0.01, Fig. 3*D*). Another key gene important for LVs is podoplanin as *Pdpn^-/-^* mice die perinatally due to failure to inflate their lungs^86^, and *Pdpn* postnatal deficiency, much akin to *Lyve1* loss, disrupts lymphangiogenesis, causes lymphedema^87^, and impedes dendritic cell migration^88^, resulting in about 20% survival^89^. We found that decorin suppressed podoplanin expression, especially in the intra-tumor LVs (Fig. 3*E*,*F*) as Lyve1. We also discovered that Hyaluronan synthase 2 (Has2), an enzyme producing hyaluronan^90–92^ was downregulated by 4-5 fold in the decorin treated allografts vis-à-vis controls (Fig. 3*G,H*). Collectively, our results demonstrate for the first time that decorin inhibits tumor progression associated lymphangiogenesis by inhibiting the RNA expression and downstream protein markers closely controlling lymphatic vessel growth. These intriguing results gave us impetus to investigate how the normal physiological process of lymphangiogenesis is affected by decorin and what implications this might have on tumor associated lymphangiogenesis.

**Fig. 3.**
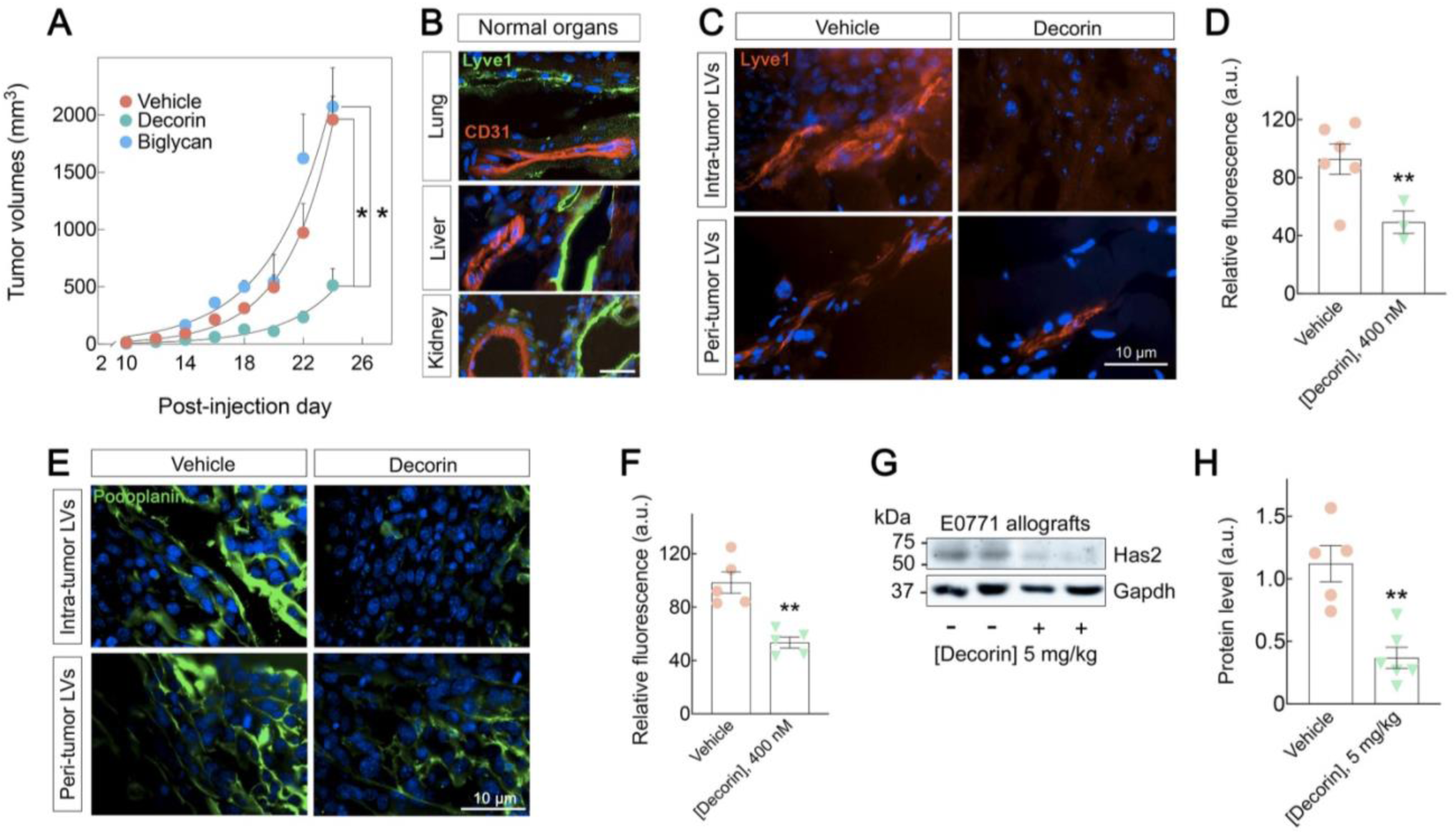
Detection of lymphatic markers in the frozen sections of breast carcinoma allografts. (A) Mammary allografts (n=5 each) ± systemic delivery recombinant decorin or biglycan, 5mg/kg, *P<0.05, one-way analysis ANOVA. (B) Distribution of Lyve1 (green) and CD31 (red) in normal mouse organs and DAPI (blue); Bar∼50 µm. (C) Distribution of Lyve1 immunostaining in the intra and peri-tumor areas, Bar∼10 µm. (D) Quantification of relative fluorescence intensity from the images captured at constant exposure time, gain and intensity; n=3-6 independent replicas, 10 fields/case;*P<0.05, unpaired two-tailed t test. (E) Immunostaining of podoplanin in intra- and peri-tumor LVs, Bar∼10 µm. (F) Quantification of relative podoplanin fluorescence, 10 fields/case, captured at identical exposure time and intensity;**P<0.01, unpaired two-tailed t test; n=5 independent biological replicates (G,H) Western blot and quantification of tumor lysates immunoreacted for Has2 and GAPDH;**P<0.01; n=5 independent biological replicates.

### Effects of Decorin in an *ex vivo* Model of Lymphangiogenesis in 3D Collagen Matrix

To address the decorin role in LV formation, we refined a 3D *ex vivo* lymphatic ring assay^93^ (Fig. S1). First, we identified thoracic ducts using Evans Blue (EB), a vital dye that is absorbed via the lymph^94^. Following EB injection in the foot pads of anesthetized mice (6-8 min), we sacrificed the mice and dissected the thoracic duct labeled by EB (Fig. 4*A*). We then sectioned the ducts into 1-mm rings and embedded them into a 3D Collagen I gel supplemented with LEC media. After four days day of lymphatic ring implantation within collagen, small branch-like outgrowth was observed emanating from the rings and with time the sproutings spread even further (Fig S2). The capillary-like structures were positive for Lyve1 by confocal microscopy (Fig. 4*B*). Notably, decorin caused a marked suppression of LV sprouts (Fig. 4*D*) vis-à-vis vehicle (Fig. 4*C*). To better quantify these findings, we used a larger cohort of mice (n=10) and found that that decorin significantly inhibited lymphatic sprouting (*P<0.001;* n=40-45 rings from 10 mice, Fig. 4 F,H) vis-à-vis vehicle (Fig. 4*E*,*H*). Sprouting area of each LRs was calculated by measuring the axial radial distance as shown in Fig S3. Importantly, equimolar amounts of recombinant biglycan did not show any inhibitory effects (Fig. 4*G*,*H*), indicating specificity for decorin bioactivity and also supporting the *in vivo* data presented in Figure 3.

**Fig. 4.**
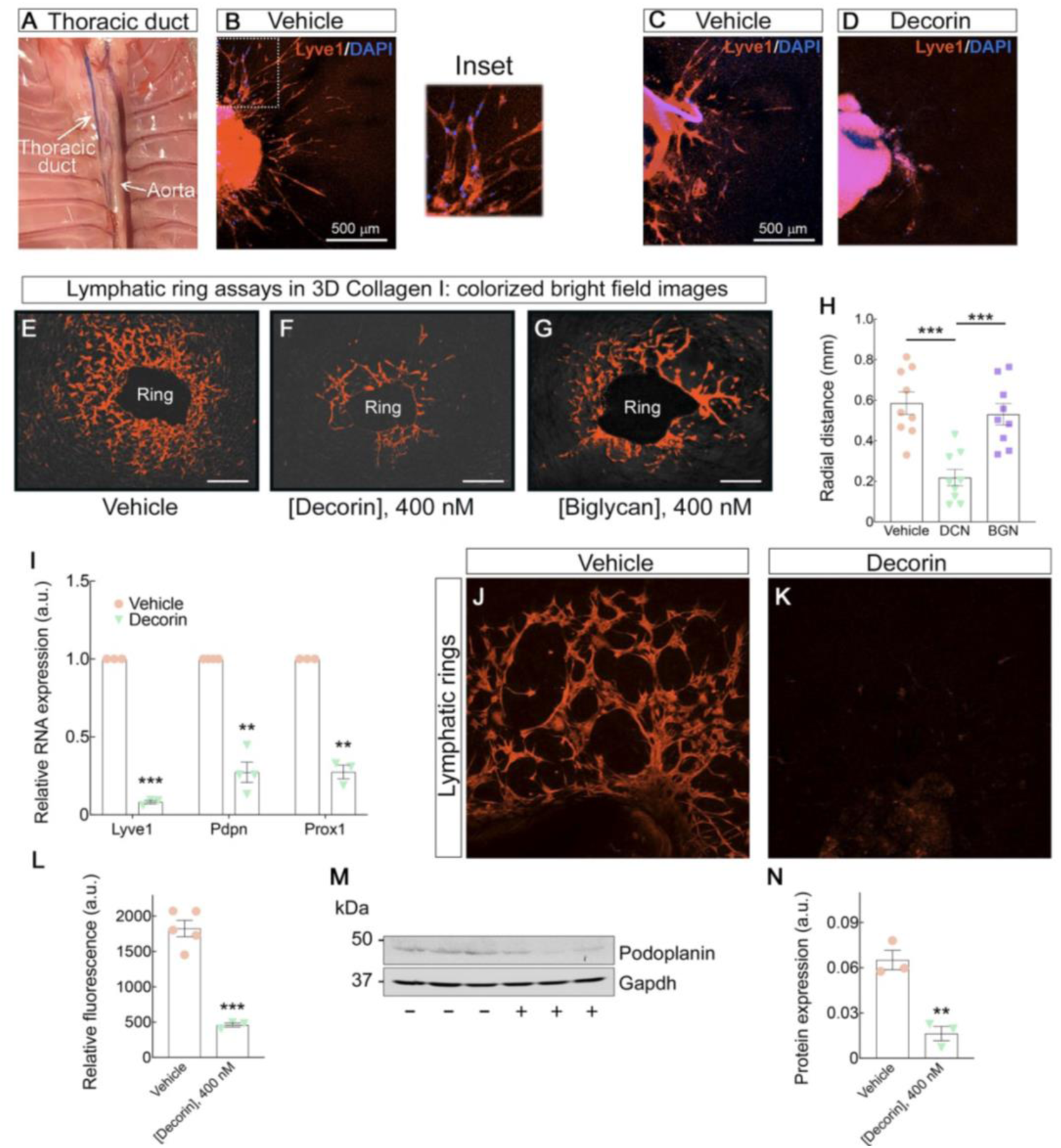
Effects of decorin in an *ex vivo* model of lymphangiogenesis in 3D Collagen matrix. (*A*) Inner view of the thorax after removal of heart and lungs. The thoracic duct has absorbed Evans blue and can be observed to the left of the thoracic aorta. (*B*) Confocal image of LR sprouts immunostained for Lyve1 (red), confirming its lymphatic origin, DAPI (blue). (*C, D*) Confocal images of Lyve1-labeled lymphatic sproutings ± decorin. (*E-G*) Colorized bright field images of lymphatic rings after day 6 of explant in 3D Collagen I. Recombinant decorin or biglycan was added for the last 3 days in culture. Bar∼400 µm. (*H*) Quantification of radial distance from five independent experiments using 10 mice, 40-45 rings/condition;\*\**P*<0.01, one way ANOVA. (*I*) qPCR analysis from 3 pooled LR sprouts ± decorin. Actin was used as a housekeeping control; mean ± SD;\*\**P*<0.01, \*\*\**P*<0.001, unpaired two-tailed *t* test, n=at least 3 biological replicates for each gene tested. (*J*) Confocal images of podoplanin (red) immunostaining in LRs ± decorin, DAPI (blue). (*K*) Quantification of mean relative fluorescence intensity captured at constant exposure time, gain and intensity; \*\*\**P*<0.001, unpaired *t* test, n= at least 3 independent biological replicas. (*L, M*) Western blot and quantification of podoplanin expression from 3 pooled LR lysates ± decorin. GAPDH used as a housekeeping control. \*\**P*<0.01, unpaired *t* test, n= 3 independent biological samples.

Next, we tested whether we could detect and quantify mRNA and proteins from the thoracic duct rings. We discovered that mRNA of three primary lymphatic markers (Lyve1, Podoplanin, and Prox-1) was markedly suppressed by chronic treatment −3 days after the initial sprouting-with recombinant decorin (Fig. 4*I*). In support of these mRNA findings, we found a four-fold reduction of Podoplanin immunoreactivity (Fig.4 *J*,*K*) which was also corroborated by immunoblotting (Fig. 4*L*,M). We also found a significant suppression of Prox1 levels (not shown). Collectively, these results provide robust evidences for a soluble proteoglycan that subdues not only tumor-associated lymphangiogenesis but also physiological lymphangiogenesis by controlling the expression of several genes that are crucial for initial lymphatic vessel development.

### Decorin Evokes Autophagic Degradation of Lyve1 in Ex Vivo Lymphatic Ring Assays and in Primary Cultures of Lymphatic Endothelial Cells

To assess whether decorin would induce a pro-autophagic program in *ex vivo* lymphatic sprouting in 3D Collagen I gels, we performed both qPCR and immunoblotting of pooled lymphatic ring lysates. We discovered that recombinant decorin under the same experimental conditions described above evoked a marked increase in mRNA for both Beclin-1 and LC3 (Fig. 5*A*). At the protein levels, we found a marked increase in both Beclin-1 (Fig. 5*B*,*C*) and LC3-II (Fig. 5*D*,*E*) in *ex vivo* LR assys. Further, using primary cultures of murine dermal LECs, we discovered that decorin evoked Beclin-1 protein levels (Fig. 5*F*,*G*). Using immunofluorescence microscopy, we found the formation of large LC3^+^ autophagosomes that persisted up to 18 h (Fig. 5*HJL*). Notably, Lyve1 co-localized with both LC3^+^ (Fig. 5K-M) and Beclin-1^+^ autophagosomes (Fig. 5N-*P*). To validate these data, we used two specific mTOR inhibitors, Torin 1^95;96^ and NK128^97^.

**Fig. 5.**
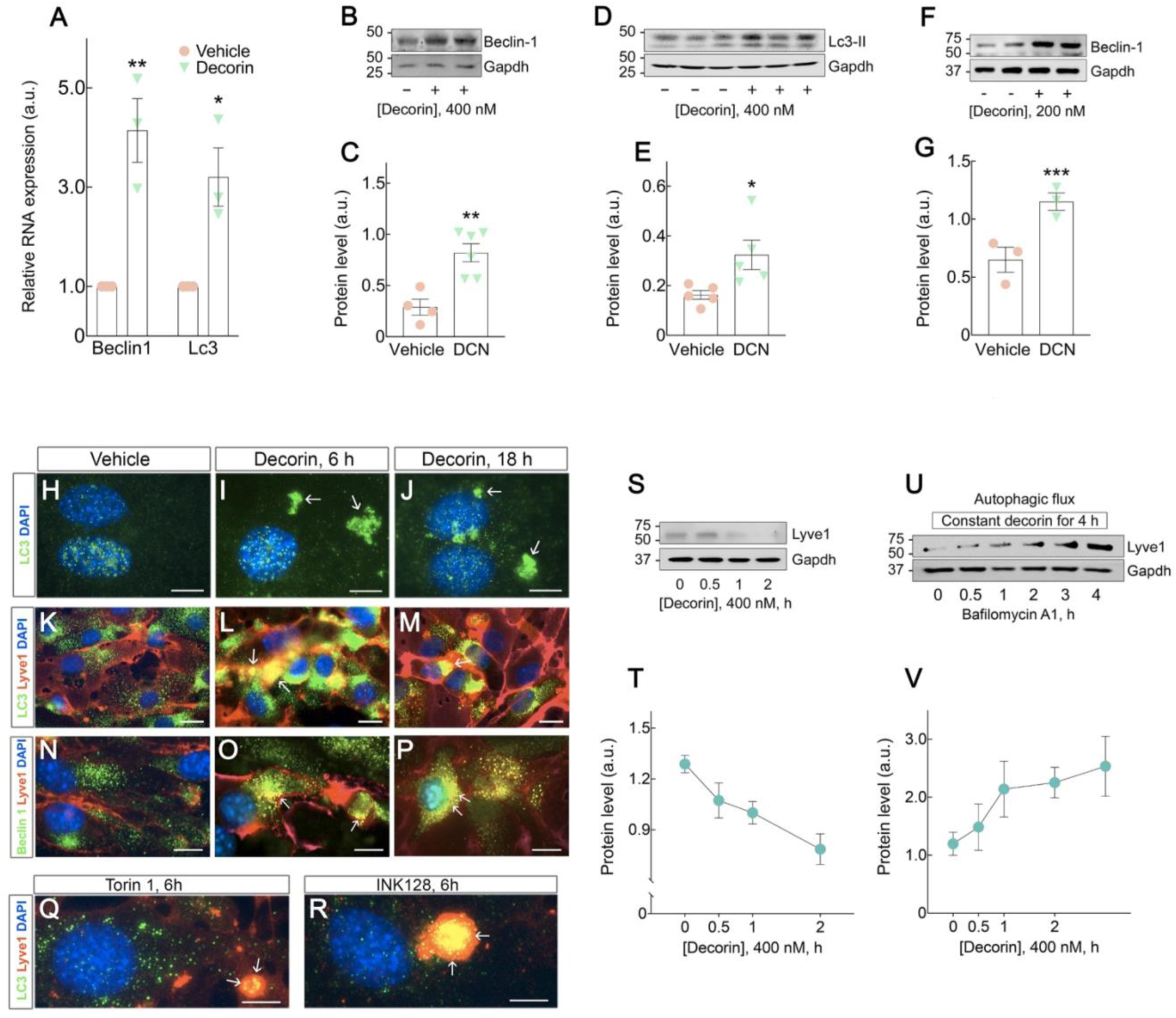
Decorin evokes autophagic degradation of Lyve1 in *ex vivo* lymphatic ring assays and in primary cultures of lymphatic endothelial cells. (*A*) qPCR analysis from 3 pooled LR sprouts ± decorin, Actin was used as an housekeeping control; mean ± SD;\*\**P*<0.01,\**P*<0.05, unpaired *t* test, n= 3 biological replicates for each gene tested. (*B-E*) Immunoblottings for LC3-II or Beclin-1 of decorin-treated LRs for 3 days and respective quantifications. GAPDH was used as housekeeping control. \*\**P*<0.01,\**P*<0.05; unpaired *t* test, n= at least 3 biological replicas for each protein. (*F,G*) Immunoblotting of primary mouse LECs for Beclin-1 and Gapdh ±decorin for 6 h and respective quantification; n=3 independent replicas; \*\*\**P* <0.001. (*H-P*) Immunodetection of LC3 and Lyve1 ±decorin (200 nM) for 6 or 18 h, as indicated; Bar∼ 10 μm. (*Q,R*) Immunodetection of LC3 and Lyve1 in LECs treated with mTOR inhibitors Torin-1 or INK128 (20 nM each) for 6 h. Arrows point to LC3+ or LC3+/Lyve1+ puncta. Bars∼10 μm. (*S,T*)Immunoblotting of primary mouse LECs for Lyve1 and Gapdh ±decorin for 6 h and respective quantification; n=3 independent replicas; \*\*\**P*<0.001. (*U,V*) Autophagic flux experiments maintaining constant decorin for 4 h ±Bafilomycin A1 (500 nM) for the indicated times. n=3 independent experiments; \**P*<0.05.

In both instances, we found similar LC3^+^/Lyve1^+^ aggregates (Fig. 5Q-RS). Furthermore, decorin downregulated Lyve1 protein levels (Fig. 5*S-T*). To confirm an activation of autophagy by decorin, we performed autophagic flux using Bafilomycin A1^98^, a V-ATPase inhibitor of autophagosome-lysosome fusion^99^. This approach measures the build-up of autophagic substrates evoked by autophagy inducers^100^. We found a progressive build-up of Lyve1 following Bafilomycin A1 treatment (Fig. 5*U*,*V*). Collectively, these results validate the imaging data presented above and provide robust evidence for decorin-evoked autophagic catabolism of Lyve1 and potentially of other key proteins of lymphatic vessels.

### Decorin Protein Core Binds and Down-regulates VEGFR3, and Activates Akt in a VEGFR3-dependent Manner in LECs

Based on the vast body of evidence implicating decorin as a pan-RTK inhibitor, we hypothesized that decorin could affect VEGFR3, the main RTK of LVs. To address this new hypothesis, we performed solid-phase binding assays using soluble Fc-tagged VEGFR3 (sVEGFR3-Fc, R&D) and decorin protein core. We have previously shown that the Fc fragment does not interfere with decorin binding to either VEGFR2^23^ or IGF-IR^22^ and that binding is wholly mediated by decorin protein core. We discovered that decorin protein core bound with relatively high affinity (Kd∼27 nM ±8) to sVEGFR3 (Fig. 6*A*) and that sVEGFR3 bound to immobilized decorin protein core with similar affinity (Kd∼47 nM ±11, Fig. 6*B*). Notably, decorin protein core (50 nM constant), was only partially displaced (∼20%) by VEGFC, even when 12 molar excess was used (Fig. 6*C*). Next, we performed dose-response experiments and found that VEGFR3 was physically downregulated by decorin within 6 h, with as little as 100 nM and slightly more reduced at higher molarities (Fig. 6*D*). These data indicate that decorin binds to the ectodomain of VEGFR3 in a region partially overlapping the natural ligand, VEGFC, causing a physical receptor downregulation. To further validate our cell system, we first demonstrated that VEGFC indeed evoked phosphorylation of Akt at Ser^473^, an established activating resi

**Fig. 6.**
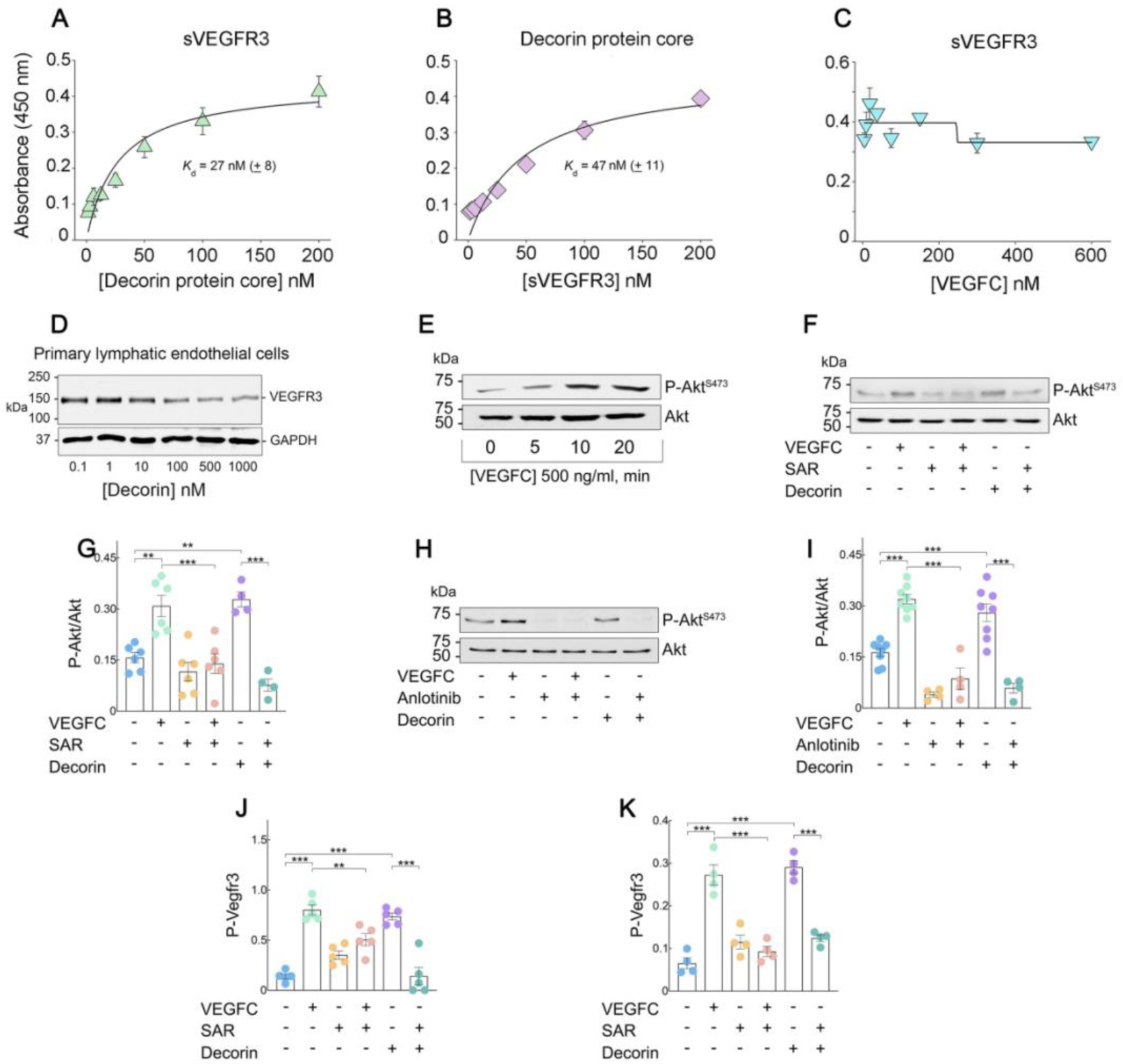
Decorin protein core binds and down-regulates VEGFR3 and activates Akt in a VEGFR3-dependent manner in LECs. (A,B) Ligand-binding assays using sVEGFR3 or decorin protein core (100 ng each) as immobilized substrates or as soluble ligands as indicated. (C) Displacement of decorin protein core (constant 50 nM) bound to immobilized sVEGFR3 by increasing concentrations of VEGFC; n=3 independent experiments run in triplicates. (D) Immunoblotting of VEGFR3 and GAPDH with increasing decorin concentrations for 6 h. (E) Immunoblotting of P-Akt induction via VEGFC. (F) Immunoblotting of P-Akt blockage evoked by the RTKi, SAR. Cells were pre-treated for 30 min with SAR 5 mM after which decorin and VEGFC were added, 400 nM and 500 ng/ml respectively, for 20 min. (G) Quantification of Akt phosphorylation at Ser473; n=4-7 biological replicates;**P<0.01;***P<0.001. (H) Immunoblotting of P-Akt blockage evoked by the RTKi, Anlotinib, and respective quantification. Cells were treated identically as in panel F.(I) Quantification of Akt phosphorylation at Ser^473^, in the presence of decorin, VEGFC or Anlotinib as indicated. n=4-7 biological replicates;**P<0.01;***P<0.001. (J) Quantification of relative P-VEGFR3 level at Tyr residues by sandwich ELISA assays in LECs treated for 20 min with either VEGFC or decorin and blocked with SAR as indicated; n= 5 biological replicates, **P<0.01, ***P<0.001. (K) Quantification of relative P-VEGFR3 level at Tyr residues by sandwich ELISA assays in LECs treated for 20 min with either VEGFC or decorin and blocked with Anlotinib as indicated; n= 5 biological replicates, ***P<0.001.

Then, we tested SAR131675 (SAR), a small molecule TKi very selective for VEGFR3 kinase^102–104^. Notably, decorin evoked phosphorylation of Akt, like VEGFC, and both activities were significantly blocked by SAR (Fig. 6*F*,*G*). To further corroborate the involvement of VEGFR3 in the decorin-evoked signaling, we tested Anlotinib, a novel pan-TKi with high selectivity for VEGFR2 and VEGFR3^105^, and known to deter lymphangiogenesis and LV metastasis through suppression of VEGFR3 phosphorylation^106^. We found a significant inhibition of P-Akt by Anlotinib with even more clear suppression of both basal and induced Akt phosphorylation (Fig. 6*H*,*I*). To test the efficacy of decorin-evoked phosphorylation of VEGFR3, we performed a sandwich ELISA assay which measures overall tyrosine phosphorylation of VEGFR3^107^ in human LECs ±SAR or ±Anlotinib. Both VEGFC and decorin induced VEGFR3 Tyr phosphorylation at similar levels after 20 min of exposure, and both activities were efficiently blocked by both SAR and Anlotinib (Fig. 6*J*). Collectively, these results provide robust evidence that decorin interacts with VEGFR3, not only physically but also functionally, and support the hypothesis that decorin-induced lymphangiostasis likely involves this RTK axis.

## DISCUSSION

Breast cancer is the most common form of cancer world-wide, with an increase in incidence rate each year. Developing new clinical strategies and therapeutic agents to curb breast cancer growth and metastasis are of utmost importance. Here, we have identified a new mechanism through which decorin could curtail breast cancer growth and induce tumor lymphangiostasis. Lymphangiogenesis becomes even more critical during tumor growth as it provides a smooth channel for the tumor cells to invade and proliferate by forming new lymphatic vessels. Therefore, blocking the generation of new lymphatic vessels and creating unfavourable conditions around the tumor stroma might be a strategiy to restrict cellular growth in the tumor lymphatics. Studies of the lymphangiogenic pathway have provided promising therapeutic targets and novel rationale for future cancer metastasis control ^108^. Through TCGA analysis and single cell RNAseq between a cohort of normal and breast carcinoma tissues from patients, surprisingly, we observed a significant reduction of DCN expression in the solid tumors, and with increase in the aggressiveness of tumor, decorin expression is even further diminished. These analyses clearly indicate that decorin acts as a suppressor of tumor growth and by reducing their own decorin expression, cancer cells venture to make a favourable microenvironment for transit through the vessels. To test whether restricted decorin expression in the breast carcinoma allografts has any correlation with tumor lymphangiogenesis, RNAseq analysis of the tumor allografts ± decorin was performed and a cluster of oncogenic genes were downregulated, including Lyve1, in presence of decorin. Although *Lyve1^-/-^* mice do not have an overt phenotype^109;110^, they exhibit a cell trafficking defect^111^. Thus, despite some genetic compensation, Lyve1 is very important for leukocyte entry and trafficking in LVs^111^. Moreover, Lyve1 ablation prevents docking and transit of leukocytes through LVs^112^. The unforeseen decorin-mediated suppression of Lyve1 provides a new paradigm for this soluble proteoglycan as s a novel lymphangiostatic factor contributing to the suppression of metastasis.

Not all the SLRPs have the same oncosuppressive effect like decorin. By utilizing recombinant biglycan, the most homologous SLRP to decorin^80^, we uncovered that curtailing of tumor growth, inhibition of tumor and physiological lymphangiogenesis were unique functions of decorin. The oncosuppressive effects of decorin were further corroborated in this study by showing the *in vivo* suppression of the oncoprotein β-catenin, a well-known target of decorin^21^, and the endothelial marker CD31^71^ in the breast carcinoma allografts. Our RNAseq results with the mammary allografts were validated by immunoblotting and imaging studies which showed that not only the levels of Lyve1 but also those of Podoplanin, an important LV marker, were repressed by systemic delivery of recombinant decorin. Podoplanin is another key gene important for LVs development that encodes for a Type I transmembrane sialomucin-like protein that is widely used as a biomarker for lymphatic endothelial cells, fibroblastic reticular cells of lymphoid organs and for lymphatics in the skin and tumor microenvironment. Notably, *Pdpn^-/-^* mice die perinatally due to failure to inflate their lungs^86^, and Pdpn postnatal deficiency, much akin to Lyve1 loss, disrupts lymphangiogenesis, causes lymphedema^87^, and impedes dendritic cell migration^88^, resulting in about 20% survival^89^. Podoplanin^+^ tumor lymphatics are rate limiting for breast cancer metastasis as suppression of lymphangiogenesis reduces metastases in various breast cancer models^113^. Moreover, high LV density predicts poor prognosis in breast cancer^114;115^. As tumor cells utilize similar mechanisms as leukocytes in migrating through LVs^116;117^, we hypothesize that Lyve1 and Podoplanin would also have an effect on cancer cell trafficking through LVs, ultimately affecting breast cancer dissemination and metastasis.

To further explore the efficacy of exogenous decorin in lymphangiogenesis we used a refined *ex vivo* 3D collagen matrix assay based on mouse thoracic duct explants. Chronic decorin treatment markedly suppressed the radial distance of the lymphatic sprouts vis-à-vis vehicle, suggesting that decorin is not only affecting tumor but also physiological lymphangiogenesis. Notably, Lyve1^+^/Podoplanin+ cells in the sprouted lymphatic rings and real time analysis of mRNA expression established that most lymphatic markers were indeed expressed by lymphatic outgrowths. Decorin treatment significantly reduced the mRNA expression of lymphatic markers like Lyve1, Podoplanin and Prox1 clearly inferring that decorin induces downregulation of several genes, important for initial lymphatic vessel development. Recently, it has been shown that in squamous cell carcinoma, cell migration and invasion is suppressed along with reduced metastasis *in vivo* by Lyve1 knockdown, ^118^. Furthermore, loss of Lyve1 has been established to protect against hepatic melanoma metastasis and a monoclonal antibody against Lyve1 has been identified in suppressing breast tumor growth and lymph node metastasis^119;120^.Similar to Lyve1, podoplanin expression in the tumor stroma has also been shown to increase epithelial mesenchymal transition, migration, invasion and induce lymphangiogenesis^121^. In lung squamous cancer cells, podoplanin was found to inhibit tumor associated lymphangiogenesis via suppression of endogenous VEGFC^122^. Collectively, these findings support the concept that Lyve1 and Podoplanin are two key biomarkers for many cancer types that might provide a powerful tool in designing strategies for future cancer treatment.

As decorin is shown to evoke endothelial cell autophagy via Peg3, we further investigated the autophagic machinery in lymphatic rings in the absence or presence of recombinant decorin and particularly attending to the existing correlation between induced lymphangiostasis and autophagy^23;123^. Increased Beclin-1 and LC3 accumulation, two crucial autophagic markers, at both mRNA and protein level, corroborated our findings of evoked autophagy in endothelial cells^23^. Consistent with this, decorin induced co-localization of Lyve1 with either Beclin1 or LC3 was validated using two specific mTOR inhibitors. Although autophagy in cancer cells is considered to be a double edged sword, decorin mediated upregulation of autophagy may be anti-tumorigenic because basal level of autophagy is important for tumor growth and metastasis, and lost or aberrant autophagy leads to induction of genomic instability and necrosis in mouse tumor models^124^. We also observed that chronic decorin exposure in LECs using the autophagic flux inhibitor Bafilomycin A1, evoked a build-up of Lyve1. Thus, the dual activity of decorin could be exploited to prevent systemic spreading of breast cancer in a clinical setting. There is strong evidence indicating that the VEGFR3 signalling axis is required for lymphangiogenesis but not for angiogenesis^125;126^.

Indeed, VEGFR3 activation in breast cancer cells promotes tumor lymphangiogenesis and metastases^127^, and VEGFR3 has been linked to AMPK-mediated autophagy^128^, further supporting our current and previous results^23;123;129^. Thus we hypothesized that soluble decorin, being a natural pan-RTK inhibitor, could interact with VEGFR3. Our prediction was validated as we discovered that decorin protein core bound with relatively high affinity to sVEGFR3 and physically downregulated the protein level of VEGFR3. Mechanistically decorin caused a transient activation of the pVEGFR3/pAKT axis similar to VEGFC, the natural ligand of VEGFR3. However, the ultimate effects of decorin were downregulation of VEGFR3 in a way similar to that produced on EGFR and Met^18;20;24^. By downregulating VEGFR3 signalling, decorin might alter lymphatic vessel development and organization, and induce lymphangiostasis in similar way as seen in cardiac samples^130^. In this case, defective VEGFR3 signalling was associated with altered cardiac LV morphology, dilated LVs leading to a higher mortality rate after acute myocardial infarction ^130^. Notably, AD0157, a pyrrolidinedione compound isolated from the fermentation broth of the marine fungus Paraconiothyrium sp. was found to mechanistically block VEGFR3 pathway in breast cancer xenografts of mice^8^. The compound was able to reduce breast tumor growth, blocks the invasion of tumor cells to the draining lymph nodes, and potently reduces metastases, through a strong reduction of the lymphatic vasculature in both primary tumors and in regional lymph nodes^8^. So, we propose here, like AD0157, decorin also has the potential to be considered as an anti-lymphangiogenic drug with the same mechanism of action. In our continuous efforts to discover and characterize new ways to tackle lymphatic metastasis, decorin mediated induction of lymphangiostasis could shed light on promising therapeutics.

### Limitations of the study

While we have thoroughly characterized the decorin-evoked downregulation of a cluster of breast cancer-associated genes involved in lymphatic development, this study has several limitations. First, the results are based on a single model of mammary carcinoma and thus our studied should be expanded to additional tumor models. Second, although we provide compelling evidence for a role of VEGFR3 in the signaling of decorin in LVs, we do not know the precise mechanism for a concurrent downregulation of Lyve1 and podoplanin at both transcriptional and post-transcriptional level. Third, though we have demonstrated a role for soluble decorin in inducing autophagic degradation of Lyve1, we do not know how many other pro-lymphangiogenic factors are catabolized via autophagy or canonical lysosomal pathway.

In conclusion, our findings reveal a previously unappreciated role of decorin in suppressing tumor lymphangiogenesis and also physiological lymphatic sprouting with implications for future therapeutic application in various forms of cancer.

## MATERIALS and METHODS

### Animals and Cells

6-8 week old C57BL/6 mice were purchased from Jackson Laboratories (Bar Harbor, ME) and housed at Thomas Jefferson University (TJU) animal research facility in sterilized filter top cages under 12-h light-dark cycle, humidity and temperature-controlled conditions. All animal procedures were approved by the Animal Care and Use Committee of TJU. E0771 cells were obtained from ATCC and were maintained in DMEM (Gibco, Thermo Fisher Scientific, Inc.) with 10% FBS (Gibco, Thermo Fisher Scientific, Inc.) and 1% penicillin-streptomycin (Corning, 30-002-CI). Mouse and human primary dermal lymphatic endothelial cells (LECs) were purchased from Cell biologics, used between passages 3 to 6, and were maintained in complete endothelial cell medium (Cell biologics, M1168 and H1168). 0.25% trypsin-EDTA was used to detach and re-culture cells to the next passage. Once the cells reached 70-80% confluency, single or combination treatments was performed according to the experimental need as mentioned.

### Orthotopic E0771 Injection in Mice, Tumor Measurement and Processing

E0771 cells were used as a syngeneic model for orthotopic breast cancer growth, spontaneous metastasis, and experimental metastasis in wild-type C57BL/6 mice^131^. To develop tumors in the breast, 1x 10^6^ E0771 cells/100 µl were orthotopically implanted into the right fat pad of the fourth mammary gland using a sterile 22-gauge needle. Before injecting the E0771 cells in mice, live cell number was counted with a haemocytometer using 0.4% trypan blue solution (Corning, 25-900-CI). Throughout the study, health status of the mice was checked every 2 days. After Day 10 post-injection when manual palpation revealed existence of a mammary tumor around the injection site, mice were randomly divided into different treatment groups. Recombinant decorin and biglycan were injected at 5 mg/kg body weight, every other day until day 20 post-injection. Tumor sizes were assessed throughout the experiment by caliper measurements of length (L) and width (W), and tumor volume (V) was calculated using the formula: V (in mm^3^) = 0.5 x L x W^2^. When the largest tumor reached 2000 mm^3^ in size, mice were sacrificed by CO_2_ inhalation and tumors were surgically dissected and divided into two halves. One half of the sample was flash frozen in liquid N_2_ and stored in −80°C for further biochemical analysis. The other half was kept in 10% buffered formalin for 24 h and transferred through a sucrose gradient. The sample is soaked in 15% sucrose for 24 h followed by 30% sucrose for another 24 h. The sample is then embedded in OCT (Sakura Finetek, CA, USA) medium and stored at −80°C. 8-10 µm thick sections were cut utilizing a cryostat (Leica) from the frozen tumor tissue, mounted on glass slides and further processed for immunofluorescence analysis.

### Preparation of 3D Collagen Type I Matrix and Thoracic Duct Isolation from Mice

We used a mixture of rat-tail collagen type I (Corning, 354236); media 199 10X phenol red free (ThermoFisher Scientific); 140 mM NaHCO_3_ (filter sterilized); 1M NaOH (dissolved in autoclaved water) and ultrapure sterile water. At neutral pH, buffered collagen mixture was added to the desired culture plate to cover the bottom and allowed to polymerize at 37°C. Following anesthetization with isoflurane, 4% Evan’s blue was injected into both hind footpads of the C57BL/6 wild type mice and blue stained dye was allowed to travel through the lymphatic system and stain thoracic duct for 6-8min. After sacrificing the mice with CO_2_ inhalation, the blue-stained thoracic duct adjacent to the aorta was carefully dissected out from the opened chested cavity as shown in Fig S1. After trimming most of the fat from the isolated thoracic duct, it was chopped into small pieces and each small piece was embedded in the previously prepared culture plate with type I collagen. Each piece of lymphatic ring was sandwiched with another layer of top collagen, and endothelial cell media (Cell Biologics, M1168) was added on top of it to facilitate lymphatic vessel sprouting at 37°C in 5% CO_2_. Here, we were successfully able to grow at least 20-25 rings of ∼1 mm width from the thoracic duct of each C57BL/6 mice. Either 400 nM of decorin or biglycan was added for 3 consecutive days after day 4 of LR explant when sproutings begin (Fig. S2). After Day 7 of growing LRs on the collagen in absence or presence 3 days of chronic decorin/biglycan, bright field images were captured at 4X in Leica DM IL LED inverted microscope for analysing through ImageJ software (https://imagej.net/Fiji/) and further processed for immunoblotting and immunostaining.

### Radial Distance of the Sprouts

Next, each captured image of the representative rings after Day 7, was analyzed using ImageJ software. We typically observed sprouting after Day 4 around the LRs. By Day 7, dense vessel sprouting could be observed from each lymphatic ring. Sprouting is mostly branched from the sides of the lymphatic ring. So, here we propose a method through ImageJ where one can precisely measure the distance travelled by the sprouts from the end of the thoracic duct. After Day 7, phase contrast images of the live explants were acquired, and using the subtract background function in ImageJ, background was removed from each image using the rolling ball radius of 700 pixels. This step reduces noise and improves contrast. Next, the sprouts were exclusively highlighted in red using the adjust threshold function, eliminating the lymphatic ring and any empty spaces. Finally, using the set scale function, the accurate pixel/micron ratio was entered based on the resolution settings of the microscope and magnification of the objective lens (i.e. 0.645 μm/pixel), and check the global setting option. Using the circle option on the ImageJ toolbar, a circle was drawn around the highlighted sprouts and another around the lymphatic ring (Fig S3). Using the measure function, the radii of each circle was determined and the radius of the larger circle was subtracted from that of the smaller one to calculate the radial distance of the sprouts.

### RNAseq, Bioinformatics Analyses and Imaging Studies

We performed high-throughput deep RNAseq (Genewiz, NJ) from the orthotopic mammary tumor allografts of 14 mice (RNAseq data available at www.iozzolab.com). We validate RNA integrity through capillary electrophoresis and mRNA was enriched and sequenced utilizing the Illumina® HiSeq® system configured at 2x150 bp. Sequence reads were trimmed to remove possible adapter sequences and poor quality nucleotides using Trimmomatic v.0.36^132^ and mapped to the Mus musculus GRCm38 reference genome (ENSEMBL) using STAR aligner v.2.5.2b^73^. Unique gene hits were calculated via Counts from the Subread v.1.5.2. At protein level, we used STRING v11 database to get score for protein/protein interactions network^133^. For immunostaining, antibodies were obtained from the following sources: anti-6x-His (1:200, ab18184), β-catenin (1:100, ab16051), CD31 (1:100, ab222783), podoplanin (1:100, ab10288), beclin1 (1:200, ab62557) from Abcam (Cambridge, MA). Lyve1 (1:50, 14-0443-82) and LC3 (1:100, L7543) were purchased from ThermoFisher Scientific (Waltham, MA) and Sigma Aldrich (St. Louis, MO). Secondary antibodies like goat anti-mouse Alexa Fluor^TM^ 488 (A-11001), goat anti-rat Alexa Fluor^TM^ 594 (A-11007), goat anti-mouse Alexa Fluor^TM^ 594 (A-11004) and donkey anti-rabbit Alexa Fluor^TM^ 488 (A-21206) were purchased from ThermoFisher Scientific and used at 1:400 dilution. Tumor sections mounted on glass slide were blocked with freshly prepared 1% BSA in PBS for 1 h followed by incubation with respective primary antibody for 1 h at room temperature. Slides were washed three times with PBS after the primary antibody incubation to remove any remaining stain and further probed with respective secondary antibody at 1:400 dilution for 1 h in the dark. To remove the excess stain, slides were washed again with PBS and allowed to dry for 1 min. VECTASHIELD antifade mounting medium with DAPI (Vector Laboratories, H-1500) was added and a clean coverslip was placed on top of the tissue section and finally sealed before visualizing in Leica DM5500 B fluorescence microscope. 3D collagen embedded LRs treated with and without 3 days of chronic decorin and mouse LECs were fixed with 4% paraformaldehyde (PFA) for 30 min followed by permeabilization with 0.1% Triton-X100. Mouse LECs were grown on a glass coverslip in a 12 well culture plate. 1% BSA in PBS was used as blocking agent for 1 h and primary antibodies diluted in blocking solution was added for overnight incubation. Next day, after washing the primary antibody with PBS by gentle shaking, respective secondary antibodies were added and incubated for 2 h. To remove the excess unbound secondary, samples were again washed three times with PBS, stained with DAPI and finally visualized in either Leica DM5500 B fluorescence or Nikon A1R inverted confocal microscope.

### Immunoblotting and Inhibitors

For Western blotting, antibodies were procured from the following sources: β-catenin (1:1000, ab16051), CD31 (1:1000, ab222783), podoplanin (1:500, ab10288) and Beclin1 (1:2000, ab62557) from Abcam. Has2 (sc-514737) was purchased from Santa Cruz Biotechnology (1:500, Santa Cruz). Lyve1 (1:500, #67538), GAPDH (1:4000, #2118), Phospho-Akt (Ser473) (1:500, #9271) and Akt antibody (1:1000, #9272) were obtained from Cell Signaling Technology (Beverly, MA). Human VEGFR3/Flt-4 antibody (1:500, AF743) was obtained from R&D systems (Minneapolis, MN). Horse Radish Peroxidase (HRP) conjugated secondary antibodies like goat anti-rabbit (Millipore, AP307P, 1:3000), goat anti-mouse (Promega, W4021, dilution 1:3000) and rabbit anti-goat (ThermoFisher Scientific, R-21459, 1:1000) were used for this study. Tumor lysates were prepared in RIPA buffer: 50 mM Tris-Cl; pH 7.5, 150 mM NaCl, 1 mM EGTA, 1 mM EDTA, 0.5% deoxycholate, 0.5% SDS, 1% Triton-X 100, 1 mM orthovanadate, 1 mM PMSF, plus protease inhibitor cocktail, (ThermoFisher,#A32961. Equal amounts of protein after lysis were loaded on SDS-PAGE. Three sprouted LRs with or without three-day chronic decorin treatment were pooled with RIPA lysis buffer to enhance the protein yield and lysed with sonication on ice. Mouse and human LECs were cultured in 12-well plates at a density of 10^5^ cells/ml and at around 70% confluency, all the experiments were done. LECs were lysed in RIPA after scrapping and all the lysates including LR lysates were mixed with 5X SDS sample buffer (250 mM Tris-HCl, pH 6.8, 10% (wt/vol) SDS, 40% (vol/vol) glycerol, 5 % β−mercaptoethanol) and boiled at 100°C for 8-10 min. Proteins were separated on 10 % or 14% SDS-PAGE and transferred to nitrocellulose membranes (#10600002, GE Healthcare Biosciences). The membranes were blocked with 1% BSA in TBS-Tween 20 (TBST) for 1 h and then incubated with respective primary antibodies diluted in blocking solution for overnight. After washing in TBST, membranes were probed with secondary antibodies for 1 h at room temperature. Finally membranes were again washed three times with TBST and submerged in SuperSignal West Pico chemiluminescent substrate for 2 min before visualizing through ImageQuant LAS 4000 chemiluminescent Image analyzer (GE Healthcare). We used a variety of inhibitors including Torin 1 (inh-tor1) and Bafilomycin A1 (tlrl-baf1) both purchased from InvivoGen (San Diego, CA). INK128 (S2811), SAR131675 (S2842) and Anlotinib (S8726) were purchased from Selleckchem (Houston, TX).

### Quantitative PCR (qPCR) and Solid-phase Binding Assays

From a pool of 3 LR lysates, total mRNA was extracted by GenElute™ Single Cell RNA Purification Kit (Sigma-Aldrich, RNB300) as per manufacturer’s protocol. 500ng of RNA was used for complementary DNA synthesis using SuperScript cDNA Synthesis Kit (Thermo Fischer Scientific, 11750150) and utilized for real time PCR using different primer sets listed in Table S1. Quality of RNA was verified by running 2% agarose gel, and qPCR was performed on the Step One Real-Time PCR system (Applied Biosystems, Foster City, CA) using SYBR Green PCR Master Mix II (Agilent Technologies). Amplicons representing target genes and the endogenous housekeeping gene, β-actin, were amplified in at least triplicate, independent reactions. Fold change determinations were made using the Comparative Ct method for expression analysis. Delta Ct (ΔCt) values represent gene expression levels to normalized β-actin for each reaction. Delta Delta Ct (ΔΔCt) values then represent the experimental cDNA (for example, 400 nM decorin treated sample) minus the corresponding gene levels of the calibrator sample (control sample). Fold changes were calculated using the double ΔCt method (2^−ΔΔCT^) ± SEM. Direct binding of decorin protein core to the soluble Fc tagged VEGFR3 (sVEGFR3-Fc, R&D systems, 349-F4), was performed using solid-phase binding assay. ELISAs were performed according to a standard protocol as shown previously^134;135^. Decorin or sVEGFR3-Fc (100 ng/well) was allowed to adhere to the 96 well plate overnight at RT in the presence of carbonate buffer, pH 9.6. Plates were washed with PBS, incubated for 2 h with serial dilutions of decorin protein core or sVEGFR3. In the quantitative competition experiment, decorin protein core was kept at constant concentration (50 nM) and incubated with increasing concentrations of VEGFC. After incubation, plates were extensively washed with PBS, blocked with 1% BSA/ in PBS, and incubated for 1 h with an antibody raised against decorin^69^ or VEGFR3 (R&D systems). HRP-conjugated secondary antibody was incubated for 1 h and the immune complexes were revealed using SIGMA-FAST ^TM^ O-phenylenediamine dihydrochloride (P9187). Absorbance at 490 nm was measured in a Victor3TM (PerkinElmer Life Sciences).

### Phospho-VEGFR3 Sandwich ELISA

Phosphorylation of VEGFR3 was estimated by sandwich ELISA as shown in previous report with minor modifications^107^. Human dermal LECs were plated on 12 well plates at a density of 1 x 10^5^cell/ml and at 60-70% confluency, LECs were serum starved for 24 h at 37°C. According to the experimental need, serum starved human LECs were pre-treated with 5µM SAR131675 or Anlotinib for 30 min, before VEGF-C or decorin stimulation for 20 minutes. The media were then aspirated and the cells washed once with cold PBS before being lysed with 150 lysis buffer :1% NP-40 Alternative, 20 mM Tris [pH 8.0], 137 mM NaCl, 10% glycerol, 2 mM EDTA, 1 mM activated sodium orthovanadate, and 10% protease inhibitors. Then lysates were utilized exactly according to the instructions in the Human Phospho-VEGFR3/Flt-4 DuoSet IC ELISA Kit (R&D systems, DYC2724-5).

### Statistical Analyses

All statistical analyses were performed using Prism 9 (GraphPad Software, Inc., San Diego, CA), and details can be found in the respective figure legends. One-way analysis of variance (ANOVA) or Student’s *t* test were performed. *P*-values smaller than 0.05 were considered statistically significant.

## ACKNOWLEDGMENTS

We would like to thank all the member of the Iozzo’s laboratory and especially Aastha Kapoor and Thomas Neill for their help in the original stages of this work. We are also indebted to Eric Londin, Valerio Izzi and Jörn Dengjel for their valuable help with bioinformatics. The original research was supported in part by National Institutes of Health Grants RO1 CA39481, RO1 CA245311 and RO3 CA270830.

## AUTHOR CONTRIBUTION

All the authors were responsible for conceptualization, drafting, and editing of the manuscript, as well as the design and generation of all the figures. R.V.I. was responsible for the final editing and proofing of the manuscript

## REFERENCES

1. N. Escobedo, G. Oliver. Lymphangiogenesis: Origin, specification, and cell fate determination. Annu.Rev.Cell Dev.Biol 2016, 32, 677.

2. S. Jalkanen, M. Salmi. Lymphatic endothelial cells of the lymph node. Nat.Rev.Immunol. 2020, 20, 566.

3. R.H. Adams, K. Alitalo. Molecular regulation of angiogenesis and lymphangiogenesis. Nat.Rev.Mol.Cell Biol. 2007, 8, 464.

4. D. Royston, D.G. Jackson. Mechanisms of lymphatic metastasis in human colorectal adenocarcinoma. J.Pathol. 2009, 217, 608.

5. M. Skobe, T. Hawighorst, D.G. Jackson, R. Prevo, L. Janes, P. Velasco, L. Riccardi, K. Alitalo, K. Claffey, M. Detmar. Induction of tumor lymphangiogenesis by VEGF-C promotes breast cancer metastasis. Nat.Med. 2001, 7, 192.

6. K.Y. Choi, G. Saravanakumar, J.H. Park, K. Park. Hyaluronic acid-based nanocarriers for intracellular targeting: interfacial interactions with proteins in cancer. Colloids Surf.B Biointerfaces. 2012, 99, 82.

7. N.E. Reticker-Flynn, W. Zhang, J.A. Belk, P.A. Basto, N.K. Escalante, G.O.W. Pilarowski, A. Bejnood, M.M. Martins, J.A. Kenkel, I.L. Linde, S. Bagchi et al. Lymph node colonization induces tumor-immune tolerance to promote distant metastasis. Cell 2022, 185, 1924.

8. M. Garcìa-Caballero, J. Paupert, S. Blacher, d. Van, V, A.R. Quesada, M.A. Medina, A. Noël. Targeting VEGFR-3/-2 signaling pathways with AD0157: a potential strategy against tumor-associated lymphangiogenesis and lymphatic metastases. J.Hematol.Oncol. 2017, 10, 122.

9. Y.W. Yeh, C.C. Cheng, S.T. Yang, C.F. Tseng, T.Y. Chang, S.Y. Tsai, E. Fu, C.P. Chiang, L.C. Liao, P.W. Tsai, Y.L. Yu et al. Targeting the VEGF-C/VEGFR3 axis suppresses Slug-mediated cancer metastasis and stemness via inhibition of KRAS/YAP1 signaling. Oncotarget. 2017, 8, 5603.

10. I. Martinez-Corral, Y. Zhang, M. Petkova, H. Ortsäter, S. Sjöberg, S.D. Castillo, P. Brouillard, L. Libbrecht, D. Saur, M. Graupera, K. Alitalo et al. Blockade of VEGF-C signaling inhibits lymphatic malformations driven by oncogenic *PIK3CA* mutation. Nat.Commun. 2020, 11, 2869.

11. N.K. Karamanos, Z. Piperigkou, A.D. Theocharis, H. Watanabe, M. Franchi, S. Baud, S. Brezillon, M. Gotte, A. Passi, D. Vigetti, S. Ricard-Blum et al. Proteoglycan chemical diversity drives multifunctional cell regulation and therapeutics. Chem.Rev. 2018, 118, 9152.

12. R.V. Iozzo, F. Chakrani, D. Perrotti, D.J. McQuillan, T. Skorski, B. Calabretta, I. Eichstetter. Cooperative action of germline mutations in decorin and p53 accelerates lymphoma tumorigenesis. Proc.Natl.Acad.Sci.USA 1999, 96, 3092.

13. X. Bi, C. Tong, A. Dokendorff, L. Banroft, L. Gallagher, G. Guzman-Hartman, R.V. Iozzo, L.H. Augenlicht, W. Yang. Genetic deficiency of decorin causes intestinal tumor formation through disruption of intestinal cell maturation. Carcinogenesis 2008, 29, 1435.

14. X. Bi, N.M. Pohl, G.R. Yang, Y. Gou, G. Guzman, A. Kajdacsy-Balla, R.V. Iozzo, W. Yang. Decorin-mediated inhibition of colorectal cancer growth and migration is associated with E-cadherin *in vitro* and in mice. Carcinogenesis 2012, 33, 326.

15. X. Bi, X. Xia, D. Fan, T. Mu, Q. Zhang, R.V. Iozzo, W. Yang. Oncogenic activin C interacts with decorin in colorectal cancer in vivo and in vitro. Mol.Carcinog. 2015, 55, 1786.

16. D.K. Moscatello, M. Santra, D.M. Mann, D.J. McQuillan, A.J. Wong, R.V. Iozzo. Decorin suppresses tumor cell growth by activating the epidermal growth factor receptor. J.Clin.Invest. 1998, 101, 406.

17. R.V. Iozzo, D. Moscatello, D.J. McQuillan, I. Eichstetter. Decorin is a biological ligand for the epidermal growth factor receptor. J.Biol.Chem. 1999, 274, 4489.

18. G. Csordás, M. Santra, C.C. Reed, I. Eichstetter, D.J. McQuillan, D. Gross, M.A. Nugent, G. Hajnóczky, R.V. Iozzo. Sustained down-regulation of the epidermal growth factor receptor by decorin. A mechanism for controlling tumor growth in vivo. J.Biol.Chem. 2000, 275, 32879.

19. M. Santra, C.C. Reed, R.V. Iozzo. Decorin binds to a narrow region of the epidermal growth factor (EGF) receptor, partially overlapping with but distinct from the EGF-binding epitope. J.Biol.Chem. 2002, 277, 35671.

20. S. Goldoni, A. Humphries, A. Nyström, S. Sattar, R.T. Owens, D.J. McQuillan, K. Ireton, R.V. Iozzo. Decorin is a novel antagonistic ligand of the Met receptor. J.Cell Biol. 2009, 185, 743.

21. S. Buraschi, N. Pal, N. Tyler-Rubinstein, R.T. Owens, T. Neill, R.V. Iozzo. Decorin antagonizes Met receptor activity and downregulates β-catenin and Myc levels. J.Biol.Chem. 2010, 285, 42075.

22. R.V. Iozzo, S. Buraschi, M. Genua, S.-Q. Xu, C.C. Solomides, S.C. Peiper, L.G. Gomella, R.T. Owens, A. Morrione. Decorin antagonizes IGF receptor I (IGF-IR) function by interfering with IGF-IR activity and attenuating downstream signaling. J.Biol.Chem. 2011, 286, 34712.

23. S. Buraschi, T. Neill, A. Goyal, C. Poluzzi, J. Smythies, R.T. Owens, L. Schaefer, A. Torres, R.V. Iozzo. Decorin causes autophagy in endothelial cells via Peg3. Proc.Natl.Acad.Sci.U.S.A. 2013, 110, E2582–E2591.

24. J.-X. Zhu, S. Goldoni, G. Bix, R.A. Owens, D. McQuillan, C.C. Reed, R.V. Iozzo. Decorin evokes protracted internalization and degradation of the EGF receptor via caveolar endocytosis. J.Biol.Chem. 2005, 280, 32468.

25. A. De Luca, M. Santra, A. Baldi, A. Giordano, R.V. Iozzo. Decorin-induced growth suppression is associated with upregulation of p21, an inhibitor of cyclin-dependent kinases. J.Biol.Chem. 1996, 271, 18961.

26. D.G. Seidler, S. Goldoni, C. Agnew, C. Cardi, M.L. Thakur, R.A. Owens, D.J. McQuillan, R.V. Iozzo. Decorin protein core inhibits *in vivo* cancer growth and metabolism by hindering epidermal growth factor receptor function and triggering apoptosis via caspase-3 activation. J.Biol.Chem. 2006, 281, 26408.

27. T. Neill, H.R. Jones, Z. Crane-Smith, R.T. Owens, L. Schaefer, R.V. Iozzo. Decorin induces rapid secretion of thrombospondin-1 in basal breast carcinoma cells via inhibition of Ras homolog gene family, member A/Rho-associated coiled-coil containing protein kinase 1. FEBS J. 2013, 280, 2353.

28. C.C. Reed, A. Waterhouse, S. Kirby, P. Kay, R.A. Owens, D.J. McQuillan, R.V. Iozzo. Decorin prevents metastatic spreading of breast cancer. Oncogene 2005, 24, 1104.

29. C.C. Reed, J. Gauldie, R.V. Iozzo. Suppression of tumorigenicity by adenovirus-mediated gene transfer of decorin. Oncogene 2002, 21, 3688.

30. S. Goldoni, D.G. Seidler, J. Heath, M. Fassan, R. Baffa, M.L. Thakur, R.A. Owens, D.J. McQuillan, R.V. Iozzo. An anti-metastatic role for decorin in breast cancer. Am.J.Pathol. 2008, 173, 844.

31. W. Xu, T. Neill, Y. Yang, Z. Hu, E. Cleveland, Y. Wu, R. Hutten, X. Xiao, S.R. Stock, D. Shevrin, K. Kaul et al. The systemic delivery of an oncolytic adenovirus expressing decorin inhibits bone metastasis in a mouse model of human prostate cancer. Gene Therapy 2015, 22, 31.

32. J.G. Tralhão, L. Schaefer, M. Micegova, C. Evaristo, E. Schönherr, S. Kayal, H. Veiga-Fernandes, C. Danel, R.V. Iozzo, H. Kresse, P. Lemarchand. In vivo selective and distant killing of cancer cells using adenovirus-mediated decorin gene transfer. FASEB J. 2003, 17, 464.

33. K. Shintani, A. Matsumine, K. Kusuzaki, J. Morikawa, T. Matsubara, T. Wakabayashi, K. Araki, H. Satonaka, H. Wakabayashi, T. Lino, A. Uchida. Decorin suppresses lung metastases of murine osteosarcoma. Oncology Reports 2008, 19, 1533.

34. K. Araki, H. Wakabayashi, K. Shintani, J. Morikawa, A. Matsumine, K. Kusuzaki, A. Sudo, A. Uchida. Decorin suppresses bone metastasis in a breast cancer cell line. Oncology 2009, 77, 92.

35. H. Zhao, H. Wang, F. Kong, W. Xu, T. Wang, F. Xiao, L. Wang, D. Huang, P. Seth, Y. Yang, H. Wang. Oncolytic adenovirus rAd. DCN inhibits breast tumor growth and lung metastasis in an immune-competent orthotopic xenograft model. Hum.Gene Ther. 2019, 30, 197.

36. X. Hu, E.S. Villodre, R. Larson, O.M. Rahal, X. Wang, Y. Gong, J. Song, S. Krishnamurthy, N.T. Ueno, D. Tripathy, W.A. Woodward et al. Decorin-mediated suppression of tumorigenesis, invasion, and metastasis in inflammatory breast cancer. Commun.Biol 2021, 4, 72.

37. C. VanOpstall, S. Perike, H. Brechka, M. Gillard, S. Lamperis, B. Zhu, R. Brown, R. Bhanvadia, D.J. Vander Griend. MEIS-mediated suppression of human prostate cancer growth and metastasis through HOXB13-dependent regulation of proteoglycans. Elife. 2020, 9, e53600.

38. G.F. Gao, J.S. Parker, S.M. Reynolds, T.C. Silva, L.B. Wang, W. Zhou, R. Akbani, M. Bailey, S. Balu, B.P. Berman, D. Brooks et al. Before and after: Comparison of legacy and harmonized TCGA genomic data Commons’ data. Cell Syst. 2019, 9, 24.

39. S.Z. Wu, D.L. Roden, C. Wang, H. Holliday, K. Harvey, A.S. Cazet, K.J. Murphy, B. Pereira, G. Al-Eryani, N. Bartonicek, R. Hou et al. Stromal cell diversity associated with immune evasion in human triple-negative breast cancer. EMBO J. 2020, 39, e104063.

40. S. Goldoni, R.V. Iozzo. Tumor microenvironment: Modulation by decorin and related molecules harboring leucine-rich tandem motifs. Int.J.Cancer 2008, 123, 2473.

41. R.V. Iozzo, R.D. Sanderson. Proteoglycans in cancer biology, tumour microenvironment and angiogenesis. J.Cell.Mol.Med. 2011, 15, 1013.

42. S. Buraschi, T. Neill, R.T. Owens, L.A. Iniguez, G. Purkins, R. Vadigepalli, B. Evans, L. Schaefer, S.C. Peiper, Z. Wang, R.V. Iozzo. Decorin protein core affects the global gene expression profile of the tumor microenvironment in a triple-negative orthotopic breast carcinoma xenograft model. PLoS ONE 2012, 7, e45559.

43. C. Desmedt, F. Piette, S. Loi, Y. Wang, F. Lallemand, B. Haibe-Kains, G. Viale, M. Delorenzi, Y. Zhang, M.S. d’Assignies, J. Bergh et al. Strong time dependence of the 76-gene prognostic signature for node-negative breast cancer patients in the TRANSBIG multicenter independent validation series. Clin.Cancer Res. 2007, 13, 3207.

44. G. Jönsson, J. Staaf, J. Vallon-Christersson, M. Ringnér, S.K. Gruvberger-Saal, L.H. Saal, K. Holm, C. Hegardt, A. Arason, R. Fagerholm, C. Persson et al. The retinoblastoma gene undergoes rearrangements in BRCA1-deficient basal-like breast cancer. Cancer Res. 2012, 72, 4028.

45. M. Schmidt, D. Böhm, T.C. von, E. Steiner, A. Puhl, H. Pilch, H.A. Lehr, J.G. Hengstler, H. Kölbl, M. Gehrmann. The humoral immune system has a key prognostic impact in node-negative breast cancer. Cancer Res. 2008, 68, 5405.

46. J. Dumas, M.A. Gargano, G.M. Dancik. shinyGEO: a web-based application for analyzing gene expression omnibus datasets. Bioinformatics. 2016, 32, 3679.

47. B. Györffy, A. Lanczky, A.C. Eklund, C. Denkert, J. Budczies, Q. Li, Z. Szallasi. An online survival analysis tool to rapidly assess the effect of 22,277 genes on breast cancer prognosis using microarray data of 1,809 patients. Breast Cancer Res.Treat. 2010, 123, 725.

48. Z. Mihály, M. Kormos, A. Lánczky, M. Dank, J. Budczies, M.A. Szász, B. Györffy. A meta-analysis of gene expression-based biomarkers predicting outcome after tamoxifen treatment in breast cancer. Breast Cancer Res.Treat. 2013, 140, 219.

49. S. Troup, C. Njue, E.V. Kliewer, M. Parisien, C. Roskelley, S. Chakravarti, P.J. Roughley, L.C. Murphy, P.H. Watson. Reduced expression of the small leucine-rich proteoglycans, lumican, and decorin is associated with poor outcome in node-negative invasive breast cancer. Clin.Cancer Res. 2003, 9, 207.

50. G. Oda, T. Sato, T. Ishikawa, H. Kawachi, T. Nakagawa, T. Kuwayama, M. Ishiguro, S. Iida, H. Uetake, K. Sugihara. Significance of stromal decorin expression during the progression of breast cancer. Oncol.Rep. 2012, 28, 2003.

51. S.J. Li, D.L. Chen, W.B. Zhang, C. Shen, G.W. Che. Prognostic value of stromal decorin expression in patients with breast cancer: a meta-analysis. J.Thorac.Dis. 2015, 7, 1939.

52. X. Li, A. Pennisi, S. Yaccoby. Role of decorin in the antimyeloma effects of osteoblasts. Blood 2008, 112, 159.

53. Y. Hu, H. Sun, R.T. Owens, J. Wu, Y.Q. Chen, I.M. Berquin, D. Perry, J.T. O’Flaherty, I.J. Edwards. Decorin suppresses prostate tumor growth through inhibition of epidermal growth factor and androgen receptor pathways. Neoplasia 2009, 11, 1042.

54. B. Bozoky, A. Savchenko, H. Guven, F. Ponten, G. Klein, L. Szekely. Decreased decorin expression in the tumor microenvironment. Cancer Med. 2014, 3, 485.

55. A. Sainio, M. Nyman, R. Lund, S. Vuorikoski, P. Boström, M. Laato, P.J. Boström, H. Järveläinen. Lack of decorin expression by human bladder cancer cells offers new tools in the therapy of urothelial malignancies. PLoS ONE 2013, 8, e76190.

56. W. Zhang, C. Zhang, W. Tian, J. Qin, J. Chen, Q. Zhang, L. Fang, J. Zheng. Efficacy of an oncolytic adenovirus driven by a chimeric promoter and armed with decorin against renal cell carcinoma. Hum.Gene Ther. 2020, 31, 651.

57. A. Reszegi, Z. Horváth, K. Karászi, E. Regós, V. Postniková, P. Tátrai, A. Kiss, Z. Schaff, I. Kovalszky, K. Baghy. The protective role of decorin in hepatic metastasis of colorectal carcinoma. Biomolecules. 2020, 10,

58. A. Reszegi, Z. Horváth, H. Fehér, B. Wichmann, P. Tátrai, I. Kovalszky, K. Baghy. Protective role of decorin in primary hepatocellular carcinoma. Front Oncol. 2020, 10, 645.

59. Z. Horvath, I. Kovalszky, A. Fullar, K. Kiss, Z. Schaff, R.V. Iozzo, K. Baghy. Decorin deficiency promotes hepatic carcinogenesis. Matrix Biol 2014, 35, 194.

60. L. Mao, J. Yang, J. Yue, Y. Chen, H. Zhou, D. Fan, Q. Zhang, S. Buraschi, R.V. Iozzo, X. Bi. Decorin deficiency promotes epithelial-mesenchymal transition and colon cancer metastasis. Matrix Biol 2021, 95, 1.

61. A. Casey, W.R. Jr. Laster, Ross G.L. Sustained enhanced growth of carcinoma EO771 in C57 black mice. Proc.Soc.Exp.Biol Med. 1951, 77, 358.

62. A. Ewens, L. Luo, E. Berleth, J. Alderfer, R. Wollman, B.B. Hafeez, P. Kanter, E. Mihich, M.J. Ehrke. Doxorubicin plus interleukin-2 chemoimmunotherapy against breast cancer in mice. Cancer Res. 2006, 66, 5419.

63. D. Dong, C. Stapleton, B. Luo, S. Xiong, W. Ye, Y. Zhang, N. Jhaveri, G. Zhu, R. Ye, Z. Liu, K.W. Bruhn et al. A critical role for GRP78/BiP in the tumor microenvironment for neovascularization during tumor growth and metastasis. Cancer Res. 2011, 71, 2848.

64. Z. Zou, S. Bellenger, K.A. Massey, A. Nicolaou, A. Geissler, C. Bidu, B. Bonnotte, A.S. Pierre, M. Minville-Walz, M. Rialland, J. Seubert et al. Inhibition of the HER2 pathway by n-3 polyunsaturated fatty acids prevents breast cancer in fat-1 transgenic mice. J.Lipid Res. 2013, 54, 3453.

65. C.N. Johnstone, Y.E. Smith, Y. Cao, A.D. Burrows, R.S. Cross, X. Ling, R.P. Redvers, J.P. Doherty, B.L. Eckhardt, A.L. Natoli, C.M. Restall et al. Functional and molecular characterisation of EO771.LMB tumours, a new C57BL/6-mouse-derived model of spontaneously metastatic mammary cancer. Dis.Model.Mech. 2015, 8, 237.

66. T. Hiraga, T. Ninomiya. Establishment and characterization of a C57BL/6 mouse model of bone metastasis of breast cancer. J.Bone Miner.Metab 2019, 37, 235.

67. I.X. Chen, V.P. Chauhan, J. Posada, M.R. Ng, M.W. Wu, P. Adstamongkonkul, P. Huang, N. Lindeman, R. Langer, R.K. Jain. Blocking CXCR 4 alleviates desmoplasia, increases T-lymphocyte infiltration, and improves immunotherapy in metastatic breast cancer. Proc.Natl.Acad.Sci.U.S.A 2019, 116, 4558.

68. K.G. Danielson, H. Baribault, D.F. Holmes, H. Graham, K.E. Kadler, R.V. Iozzo. Targeted disruption of decorin leads to abnormal collagen fibril morphology and skin fragility. J.Cell Biol. 1997, 136, 729.

69. S. Goldoni, R.T. Owens, D.J. McQuillan, Z. Shriver, R. Sasisekharan, D.E. Birk, S. Campbell, R.V. Iozzo. Biologically active decorin is a monomer in solution. J.Biol.Chem. 2004, 279, 6606.

70. G. Candiano, M. Bruschi, L. Musante, L. Santucci, G.M. Ghiggeri, B. Carnemolla, P. Orecchia, L. Zardi, P.G. Righetti. Blue silver: a very sensitive colloidal Coomassie G-250 staining for proteome analysis. Electrophoresis 2004, 25, 1327.

71. P. Lertkiatmongkol, D. Liao, H. Mei, Y. Hu, P.J. Newman. Endothelial functions of platelet/endothelial cell adhesion molecule-1 (CD31). Curr.Opin.Hematol. 2016, 23, 253.

72. S.M. Albelda, W.A. Muller, C.A. Buck, P.J. Newman. Molecular and cellular properties of PECAM-1 (endoCAM/CD31): a novel vascular cell-cell adhesion molecule. J.Cell Biol 1991, 114, 1059.

73. A. Dobin, C.A. Davis, F. Schlesinger, J. Drenkow, C. Zaleski, S. Jha, P. Batut, M. Chaisson, T.R. Gingeras. STAR: ultrafast universal RNA-seq aligner. Bioinformatics. 2013, 29, 15.

74. H. Järveläinen, P. Puolakkainen, S. Pakkanen, E.L. Brown, M. Höök, R.V. Iozzo, H. Sage, T.N. Wight. A role for decorin in cutaneous wound healing and angiogenesis. Wound Rep.Reg. 2006, 14, 443.

75. K.N. Sulochana, H. Fan, S. Jois, V. Subramanian, F. Sun, R.M. Kini, R. Ge. Peptides derived from human decorin leucine-rich repeat 5 inhibit angiogenesis. J.Biol.Chem. 2005, 280, 27935.

76. H. Järveläinen, A. Sainio, T.N. Wight. Pivotal role for decorin in angiogenesis. Matrix Biol. 2015, 43, 15.

77. T. Neill, H. Painter, S. Buraschi, R.T. Owens, M.P. Lisanti, L. Schaefer, R.V. Iozzo. Decorin antagonizes the angiogenic network. Concurrent inhibition of Met, hypoxia inducible factor-1α and vascular endothelial growth factor A and induction of thrombospondin-1 and TIMP3. J.Biol.Chem. 2012, 287, 5492.

78. P.K. Balne, S. Gupta, J. Zhang, D. Bristow, M. Faubion, S.D. Heil, P.R. Sinha, S.L. Green, R.V. Iozzo, R.R. Mohan. The functional role of decorin in corneal neovascularization in vivo. Exp.Eye Res. 2021, 207, 108610.

79. T. Neill, C.G. Chen, S. Buraschi, R.V. Iozzo. Catabolic degradation of endothelial VEGFA via autophagy. J.Biol Chem. 2020, 295, 6064.

80. R.V. Iozzo. The family of the small leucine-rich proteoglycans: key regulators of matrix assembly and cellular growth. Crit.Rev.Biochem.Mol.Biol. 1997, 32, 141.

81. S. Banerji, J. Ni, S.X. Wang, S. Clasper, J. Su, R. Tammi, M. Jones, D.G. Jackson. LYVE-1, a new homologue of the CD44 glycoprotein, is a lymph-specific receptor for hyaluronan. J.Cell Biol 1999, 144, 789.

82. A. Aruffo, I. Stamenkovic, M. Melnick, C.B. Underhill, B. Seed. CD44 is the principal cell surface receptor for hyaluronate. Cell 1990, 61, 1303.

83. P.M. Witschen, T.S. Chaffee, N.J. Brady, D.N. Huggins, T.P. Knutson, R.S. LaRue, S.A. Munro, L. Tiegs, J.B. McCarthy, A.C. Nelson, K.L. Schwertfeger. Tumor cell associated hyaluronan-CD44 signaling promotes pro-tumor inflammation in breast cancer. Cancers.(Basel) 2020, 12,

84. R. Prevo, S. Banerji, D.J. Ferguson, S. Clasper, D.G. Jackson. Mouse LYVE-1 is an endocytic receptor for hyaluronan in lymphatic endothelium. J.Biol Chem. 2001, 276, 19420.

85. L.A. Johnson, R. Prevo, S. Clasper, D.G. Jackson. Inflammation-induced uptake and degradation of the lymphatic endothelial hyaluronan receptor LYVE-1. J.Biol Chem. 2007, 282, 33671.

86. M.I. Ramirez, G. Millien, A. Hinds, Y. Cao, D.C. Seldin, M.C. Williams. T1α, a lung type I cell differentiation gene, is required for normal lung cell proliferation and alveolus formation at birth. Dev.Biol 2003, 256, 61.

87. V. Schacht, M.I. Ramirez, Y.K. Hong, S. Hirakawa, D. Feng, N. Harvey, M. Williams, A.M. Dvorak, H.F. Dvorak, G. Oliver, M. Detmar. T1α/podoplanin deficiency disrupts normal lymphatic vasculature formation and causes lymphedema. EMBO J. 2003, 22, 3546.

88. R. Bianchi, E. Russo, S.B. Bachmann, S.T. Proulx, M. Sesartic, N. Smaadahl, S.P. Watson, C.D. Buckley, C. Halin, M. Detmar. Postnatal deletion of podoplanin in lymphatic endothelium results in blood filling of the lymphatic system and impairs dendritic cell migration to lymph nodes. Arterioscler.Thromb.Vasc.Biol 2017, 37, 108.

89. J.L. Astarita, S.E. Acton, S.J. Turley. Podoplanin: emerging functions in development, the immune system, and cancer. Front Immunol. 2012, 3, 283.

90. A. Passi, D. Vigetti, S. Buraschi, R.V. Iozzo. Dissecting the role of hyaluronan synthases in the tumor microenvironment. FEBS J. 2019, 286, 2937.

91. C.G. Chen, M.A. Gubbiotti, A. Kapoor, X. Han, Y. Yu, R.J. Linhardt, R.V. Iozzo. Autophagic degradation of HAS2 in endothelial cells: A novel mechanism to regulate angiogenesis. Matrix Biol 2020, 90, 1.

92. C.G. Chen, R.V. Iozzo. Angiostatic cues from the matrix: endothelial cell autophagy meets hyaluronan biology. J.Biol Chem. 2020, 295, 16797.

93. F. Bruyère, L. Melen-Lamalle, S. Blacher, G. Roland, M. Thiry, L. Moons, F. Frankenne, P. Carmeliet, K. Alitalo, C. Libert, J.P. Sleeman et al. Modeling lymphangiogenesis in a three-dimensional culture system. Nat.Methods 2008, 5, 431.

94. M.I. Harrell, B.M. Iritani, A. Ruddell. Lymph node mapping in the mouse. J.Immunol.Methods 2008, 332, 170.

95. C.C. Thoreen, S.A. Kang, J.W. Chang, Q. Liu, J. Zhang, Y. Gao, L.J. Reichling, T. Sim, D.M. Sabatini, N.S. Gray. An ATP-competitive mammalian target of rapamycin inhibitor reveals rapamycin-resistant functions of mTORC1. J.Biol.Chem. 2009, 284, 8023.

96. Q. Liu, C. Thoreen, J. Wang, D. Sabatini, N.S. Gray. mTOR mediated anti-cancer drug discovery. Drug Discov.Today Ther.Strateg. 2009, 6, 47.

97. A.C. Hsieh, Y. Liu, M.P. Edlind, N.T. Ingolia, M.R. Janes, A. Sher, E.Y. Shi, C.R. Stumpf, C. Christensen, M.J. Bonham, S. Wang et al. The translational landscape of mTOR signalling steers cancer initiation and metastasis. Nature 2012, 485, 55.

98. E.J. Bowman, A. Siebers, K. Altendorf. Bafilomycins: a class of inhibitors of membrane ATPases from microorganisms, animal cells, and plant cells. Proc.Natl.Acad.Sci.U.S.A 1988, 85, 7972.

99. T. Yoshimori, A. Yamamoto, Y. Moriyama, M. Futai, Y. Tashiro. Bafilomycin A1, a specific inhibitor of vacuolar -type H(+)-ATPase, inhibits acidification and protein degradation in lysosomes of cultured cells. J.Biol Chem. 1991, 266, 17707.

100. D.J. Klionsky, K. Abdelmohsen, A. Abe, M.J. Abedin, H. Abeliovich, A.A. Acevedo, H. Adachi, C.M. Adams, P.D. Adams, K. Adeli, P.J. Adhihetty et al. Guidelines for the use and interpretation of assays for monitoring autophagy (3rd edition). Autophagy. 2016, 12, 1.

101. M. Sáinz-Jaspeado, L. Claesson-Welsh. Cytokines regulating lymphangiogenesis. Curr.Opin.Immunol. 2018, 53, 58.

102. A. Alam, I. Blanc, G. Gueguen-Dorbes, O. Duclos, J. Bonnin, P. Barron, M.C. Laplace, G. Morin, F. Gaujarengues, F. Dol, J.P. Hérault et al. SAR131675, a potent and selective VEGFR-3-TK inhibitor with antilymphangiogenic, antitumoral, and antimetastatic activities. Mol.Cancer Ther. 2012, 11, 1637.

103. S.D. Hwang, J.H. Song, Y. Kim, J.H. Lim, M.Y. Kim, E.N. Kim, Y.A. Hong, S. Chung, B.S. Choi, Y.S. Kim, C.W. Park. Inhibition of lymphatic proliferation by the selective VEGFR-3 inhibitor SAR131675 ameliorates diabetic nephropathy in db/db mice. Cell Death.Dis. 2019, 10, 219.

104. M. Nihei, T. Okazaki, S. Ebihara, M. Kobayashi, K. Niu, P. Gui, T. Tamai, T. Nukiwa, M. Yamaya, T. Kikuchi, R. Nagatomi et al. Chronic inflammation, lymphangiogenesis, and effect of an anti-VEGFR therapy in a mouse model and in human patients with aspiration pneumonia. J.Pathol. 2015, 235, 632.

105. G. Shen, F. Zheng, D. Ren, F. Du, Q. Dong, Z. Wang, F. Zhao, R. Ahmad, J. Zhao. Anlotinib: a novel multi-targeting tyrosine kinase inhibitor in clinical development. J.Hematol.Oncol. 2018, 11, 120.

106. T. Qin, Z. Liu, J. Wang, J. Xia, S. Liu, Y. Jia, H. Liu, K. Li. Anlotinib suppresses lymphangiogenesis and lymphatic metastasis in lung adenocarcinoma through a process potentially involving VEGFR-3 signaling. Cancer Biol Med. 2020, 17, 753.

107. H.M. Bui, D. Enis, M.R. Robciuc, H.J. Nurmi, J. Cohen, M. Chen, Y. Yang, V. Dhillon, K. Johnson, H. Zhang, R. Kirkpatrick et al. Proteolytic activation defines distinct lymphangiogenic mechanisms for VEGFC and VEGFD. J.Clin.Invest 2016, 126, 2167.

108. S.A. Stacker, S.P. Williams, T. Karnezis, R. Shayan, S.B. Fox, M.G. Achen. Lymphangiogenesis and lymphatic vessel remodelling in cancer. Nat.Rev.Cancer 2014, 14, 159.

109. N.W. Gale, R. Prevo, J. Espinosa, D.J. Ferguson, M.G. Dominguez, G.D. Yancopoulos, G. Thurston, D.G. Jackson. Normal lymphatic development and function in mice deficient for the lymphatic hyaluronan receptor LYVE-1. Mol.Cell Biol 2007, 27, 595.

110. M.X. Luong, J. Tam, Q. Lin, J. Hagendoorn, K.J. Moore, T.P. Padera, B. Seed, D. Fukumura, R. Kucherlapati, R.K. Jain. Lack of lymphatic vessel phenotype in LYVE-1/CD44 double knockout mice. J.Cell Physiol 2009, 219, 430.

111. L.A. Johnson, S. Banerji, W. Lawrance, U. Gileadi, G. Prota, K.A. Holder, Y.M. Roshorm, T. Hanke, V. Cerundolo, N.W. Gale, D. G. Jackson. Dendritic cells enter lymph vessels by hyaluronan-mediated docking to the endothelial receptor LYVE-1. Nat.Immunol. 2017, 18, 762.

112. J.M. Vieira, S. Norman, C.C. Villa Del, T.J. Cahill, D.N. Barnette, M. Gunadasa-Rohling, L.A. Johnson, D.R. Greaves, C.A. Carr, D.G. Jackson, P.R. Riley. The cardiac lymphatic system stimulates resolution of inflammation following myocardial infarction. J.Clin.Invest 2018, 128, 3402.

113. Y. Chen, D. Keskin, H. Sugimoto, K. Kanasaki, P.E. Phillips, L. Bizarro, A. Sharpe, V.S. LeBleu, R. Kalluri. Podoplanin+ tumor lymphatics are rate limiting for breast cancer metastasis. PLoS.Biol 2018, 16, e2005907.

114. S. Zhang, D. Zhang, M. Gong, L. Wen, C. Liao, L. Zou. High lymphatic vessel density and presence of lymphovascular invasion both predict poor prognosis in breast cancer. BMC.Cancer 2017, 17, 335.

115. J. Wang, Y. Guo, B. Wang, J. Bi, K. Li, X. Liang, H. Chu, H. Jiang. Lymphatic microvessel density and vascular endothelial growth factor-C and -D as prognostic factors in breast cancer: a systematic review and meta-analysis of the literature. Mol.Biol Rep. 2012, 39, 11153.

116. S. Karaman, M. Detmar. Mechanisms of lymphatic metastasis. J.Clin.Invest 2014, 124, 922.

117. D.G. Jackson. Hyaluronan in the lymphatics: The key role of the hyaluronan receptor LYVE-1 in leucocyte trafficking. Matrix Biol 2019, 78-79, 219.

118. S. Karinen, K. Juurikka, R. Hujanen, W. Wahbi, E. Hadler-Olsen, G. Svineng, K.K. Eklund, T. Salo, P. Äström, A. Salem. Tumour cells express functional lymphatic endothelium-specific hyaluronan receptor in vitro and in vivo: Lymphatic mimicry promotes oral oncogenesis? Oncogenesis 2021, 10, 23.

119. A.S. Jauch, S.A. Wohlfeil, C. Weller, B. Dietsch, V. Häfele, A. Stojanovic, M. Kittel, H. Nolte, A. Cerwenka, M. Neumaier, K. Schledzewski et al. Lyve-1 deficiency enhances the hepatic immune microenvironment entailing altered susceptibility to melanoma liver metastasis. Cancer Cell Int. 2022, 22, 398.

120. Y. Hara, R. Torii, S. Ueda, E. Kurimoto, E. Ueda, H. Okura, Y. Tatano, H. Yagi, Y. Ohno, T. Tanaka, K. Masuko et al. Inhibition of tumor formation and metastasis by a monoclonal antibody against lymphatic vessel endothelial hyaluronan receptor 1. Cancer Sci. 2018, 109, 3171.

121. H. Krishnan, J. Rayes, T. Miyashita, G. Ishii, E.P. Retzbach, S.A. Sheehan, A. Takemoto, Y.W. Chang, K. Yoneda, J. Asai, L. Jensen et al. Podoplanin: An emerging cancer biomarker and therapeutic target. Cancer Sci. 2018, 109, 1292.

122. H. Suzuki, M. Onimaru, Y. Yonemitsu, Y. Maehara, S. Nakamura, K. Sueishi. Podoplanin in cancer cells is experimentally able to attenuate prolymphangiogenic and lymphogenous metastatic potentials of lung squamoid cancer cells. Mol.Cancer 2010, 9, 287.

123. A. Torres, M.A. Gubbiotti, R.V. Iozzo. Decorin-inducible Peg3 evokes Beclin 1-mediated autophagy and Thrombospondin 1-mediated angiostasis. J.Biol Chem. 2017, 292, 5055.

124. R. Chavez-Dominguez, M. Perez-Medina, J.S. Lopez-Gonzalez, M. Galicia-Velasco, D. Aguilar-Cazares. The double-edge sword of autophagy in cancer: From tumor suppression to pro-tumor activity. Front Oncol. 2020, 10, 578418.

125. L. Zhang, F. Zhou, W. Han, B. Shen, J. Luo, M. Shibuya, Y. He. VEGFR-3 ligand-binding and kinase activity are required for lymphangiogenesis but not for angiogenesis. Cell Res. 2010, 20, 1319.

126. E.A. Korhonen, A. Murtomäki, S.K. Jha, A. Anisimov, A. Pink, Y. Zhang, S. Stritt, I. Liaqat, L. Stanczuk, L. Alderfer, Z. Sun et al. Lymphangiogenesis requires Ang2/Tie/PI3K signaling for VEGFR3 cell-surface expression. J.Clin.Invest 2022, 132, e155478.

127. M. Skobe, T. Hawighorst, D.G. Jackson, R. Prevo, L. Janes, P. Velasco, L. Riccardi, K. Alitalo, K. Claffey, M. Detmar. Induction of tumor lymphangiogenesis by VEGF-C promotes breast cancer metastasis. Nat.Med. 2001, 7, 192.

128. L. Ma, W. Li, Y. Zhang, L. Qi, Q. Zhao, N. Li, Y. Lu, L. Zhang, F. Zhou, Y. Wu, Y. He et al. FLT4/VEGFR3 activates AMPK to coordinate glycometabolic reprogramming with autophagy and inflammasome activation for bacterial elimination. Autophagy. 20211.

129. A. Goyal, T. Neill, R.T. Owens, L. Schaefer, R.V. Iozzo. Decorin activates AMPK, an energy sensor kinase, to induce autophagy in endothelial cells. Matrix Biol. 2014, 34, 46.

130. T. Vuorio, E. Ylä-Herttuala, J.P. Laakkonen, S. Laidinen, T. Liimatainen, S. Ylä-Herttuala. Downregulation of VEGFR3 signaling alters cardiac lymphatic vessel organization and leads to a higher mortality after acute myocardial infarction. Sci.Rep. 2018, 8, 16709.

131. A. Ewens, E. Mihich, M.J. Ehrke. Distant metastasis from subcutaneously grown E0771 medullary breast adenocarcinoma. AntiCancer Res. 2005, 25, 3905.

132. A.M. Bolger, M. Lohse, B. Usadel. Trimmomatic: a flexible trimmer for Illumina sequence data. Bioinformatics. 2014, 30, 2114.

133. D. Szklarczyk, A.L. Gable, D. Lyon, A. Junge, S. Wyder, J. Huerta-Cepas, M. Simonovic, N.T. Doncheva, J.H. Morris, P. Bork, L.J. Jensen et al. STRING v11: protein-protein association networks with increased coverage, supporting functional discovery in genome-wide experimental datasets. Nucleic Acids Res. 2019, 47, D607–D613.

134. T. Neill, S. Buraschi, A. Goyal, C. Sharpe, E. Natkanski, L. Schaefer, A. Morrione, R.V. Iozzo. EphA2 is a functional receptor for the growth factor progranulin. J.Cell Biol 2016, 215, 687.

135. A. Goyal, N. Pal, M. Concannon, M. Paulk, M. Doran, C. Poluzzi, K. Sekiguchi, J.M. Whitelock, T. Neill, R.V. Iozzo. Endorepellin, the angiostatic module of perlecan, interacts with both the α2β1 integrin and vascular endothelial growth factor receptor 2 (VEGFR2). J.Biol.Chem. 2011, 286, 25947.

