## Supplementary material for "Decorin suppresses tumor lymphangiogenesis: A mechanism to curtail cancer progression": Supplemetanl Figures and Table

### SUPPLEMENTARY MATERIALS

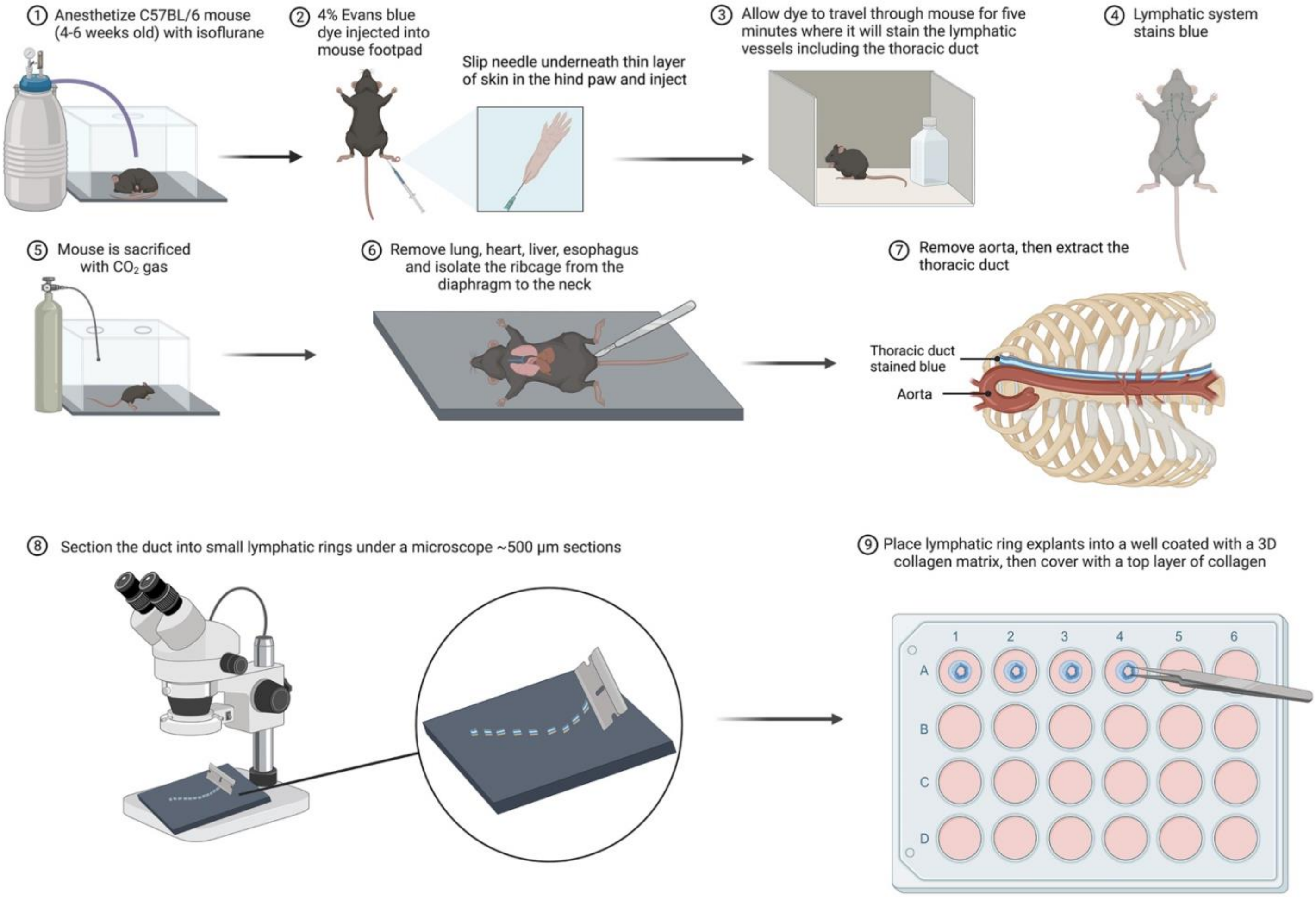


***Fig. S1***. *Schematic representation of the generation of lymphatic ring explants from mice and their growth in an ex vivo 3D matrix composed rat tail collagen type I. Every steps starting from anesthetizing the mice, removing the thoracic duct from chest cavity and growing a piece of sectioned duct as LR on 3D collagen are self-explanatory. The image was created using Biorender.com.*

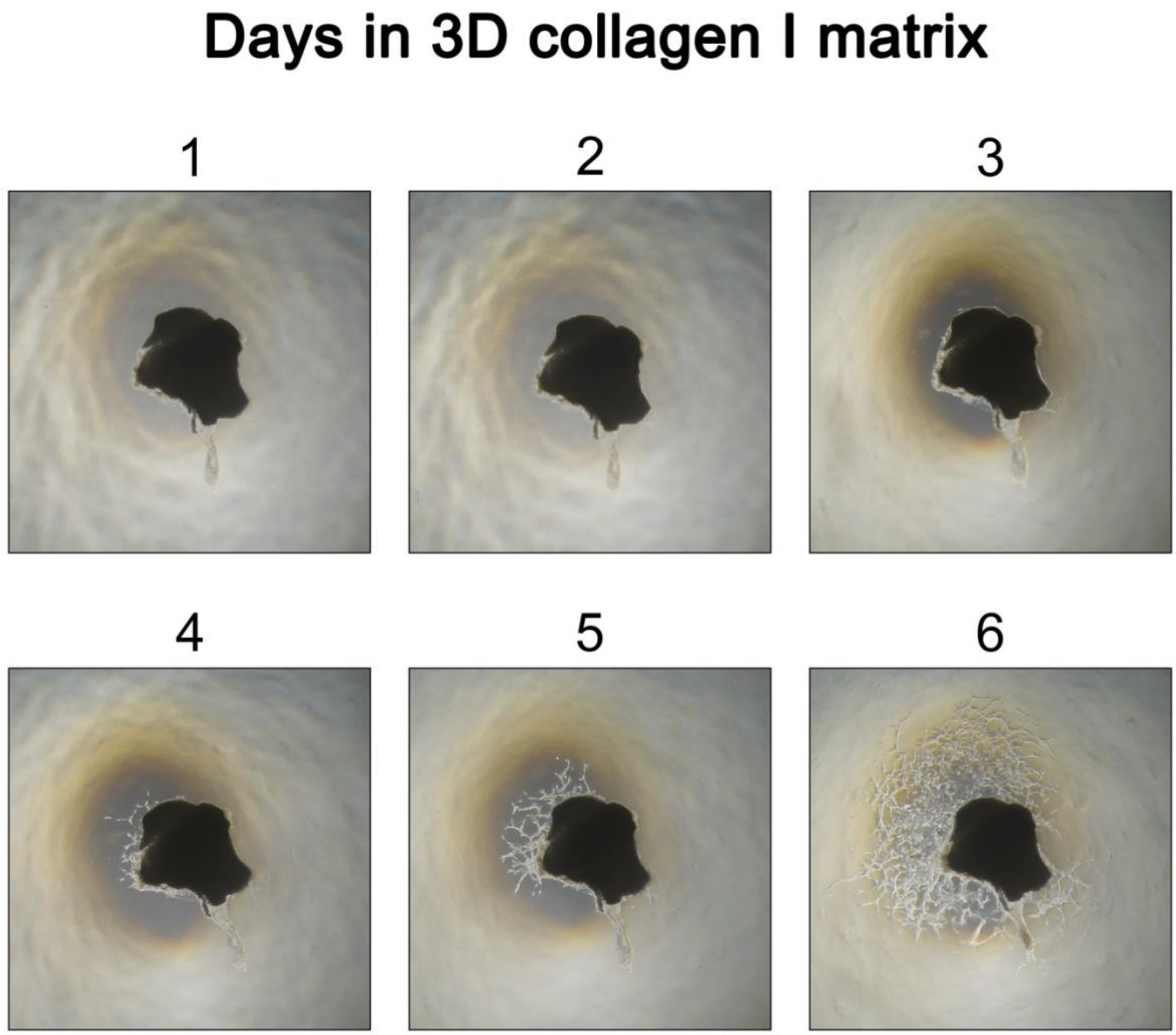


**Fig. S2.** Time course progression of lymphatic ring sprouting in the *ex vivo* assay. Bright field images of a representative lymphatic ring were taken at every day for six days at 4X magnification in an inverted microscope.

.

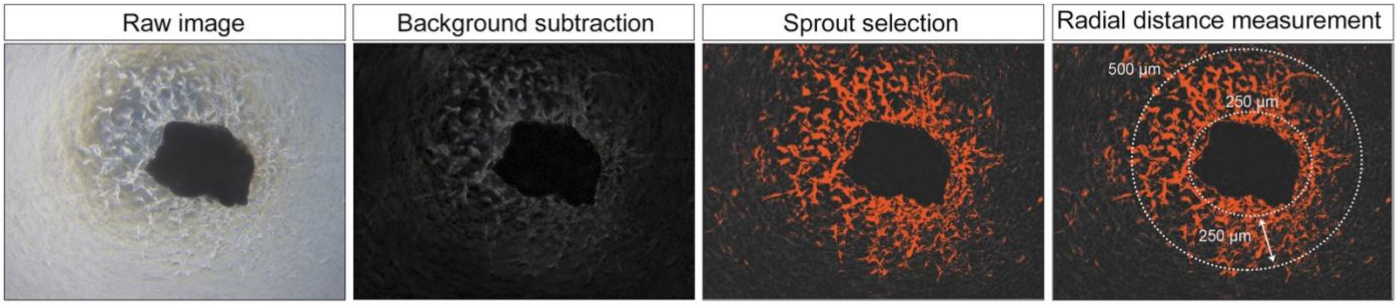


**Fig. S3.** Axial radial distance measurement of the lymphatic ring sprouting. Every steps of radial distance calculation were performed using ImageJ from the raw images, after subtracting the background and selecting only the sprouts in red.

### Supplementary table 1. List of primers used in qPCR

| Gene | Forward primer (5' – 3') | Reverse primer (3' – 5') |
|---|---|---|
| *Actb* | ACCTTCTACAATGAGCTGCG | CTGGATGGCTACGTACATGG |
| *Becn1* | TTTTCTGGACTGTGTGCAGC | GCTTTTGTCCACTGCTCCTC |
| *Cxcl12* | CAGCTCTCCTACCCTGTATCT | TGCCACAGGACAAACAGTAG |
| *Fgf7* | AAGACTGTTCTGTCGCACC | CACTTTCCACCCCTTTGATTG |
| *Fgf9* | GGAACCAGGAAAGACCACAG | TTTTCTGATCCATACAGCTCCC |
| *Map1lc3b* | TTCTTCCTCCTGGTGAATGG | GTGGGTGCCTACGTTCTCAT |
| *Lyve1* | CAGCATTCAAGAACGAAGCAG | GCCTTCACATACCTTTTCACG |
| *Mmp3* | GATGAACGATGGACAGAGGATG | AAACGGGACAAGTCTGTGG |
| *Pdpn* | GTGACCCCAGGTACAGGAGA | GCTGAGGTGGACAGTTCCTC |
| *Prox1* | CTGCTAGTGGACCTGATCTTTAC | CCTGGTCAGCTGCTTTAAGATA |
